# Environmental Variation Promotes Convergent Evolution and Rapid Diversification of Wing Shape and Color in Skipper Butterflies

**DOI:** 10.64898/2025.12.27.696669

**Authors:** Pável Matos-Maraví, Daniel Linke, Pedro Ribeiro, Gerardo Lamas, André V.L. Freitas, Rayner Núñez, Akito Y. Kawahara, Emmanuel F.A. Toussaint, Christine D. Bacon, Niklas Wahlberg, Alexandre Antonelli

## Abstract

Predation is a key driver of speciation and phenotypic diversification, yet how antipredator traits evolve and persist over evolutionary time remains poorly understood. We generated target sequence capture and whole-genome sequencing data for the skipper butterfly subfamily Eudaminae (Hesperiidae) to test whether antipredator defenses, in interaction with environmental variation, promote diversification. We focus on two antipredation wing traits: hindwing tails, which deflect predator attacks, and blue-green coloration, which may enhance motion dazzle and be used as warning coloration. Applying phylogenomics, morphometrics, and comparative methods, we model trait evolution in relation to diel activity and geographic distribution and find that hindwing tails repeatedly evolved at least seven times and blue-green coloration at least fifteen times. Both traits are associated with elevated speciation rates, but evolutionary transitions toward tailless wings and non-iridescent coloration occurred more frequently than trait gains, indicating high evolutionary lability of these antipredator defenses. Trait loss, particularly pronounced in tropical diurnal species, may reflect trade-offs in flight performance, shifts in predation guilds, or the evolution of alternative defensive traits. Our findings highlight rampant convergent evolution of wing traits that are under strong predator-mediated selection. By identifying how antipredator defenses and environmental contexts influence phenotypic and species diversification, this study provides new insights into the ecological and evolutionary processes underlying insect diversity.

---

A major question in evolutionary biology is how ecological and microevolutionary processes shape trait diversification and speciation rates (Sepkoski 1978). Predation has long been proposed as a driver of phenotypic diversification (Doebeli and Dieckmann 2000; Vamosi 2005; Ruxton et al. 2018) and speciation in prey (Rueffler et al. 2006). However, empirical evidence for predation-driven evolutionary radiations remains limited (Meyer and Kassen 2007; Pontarp and Petchey 2018), and molecular phylogenies and the fossil record sometimes disagree in corroborating or rejecting this hypothesis (Huntley and Kowalewski 2007; Arbuckle and Speed 2015; López-Villalta 2016; Zeng and Wiens 2021; Blanchard and Moreau 2023). Thus, the extent to which predation influences rates of phenotypic evolution and speciation remains a knowledge gap requiring investigation across diverse prey lineages.

Lepidoptera (butterflies and moths) provide an ideal system for determining the macroevolution of antipredator defense (Barber et al. 2015; Rubin et al. 2018, 2025; Kunte et al. 2021; Nokelainen et al. 2024). Butterfly wing shapes and color patterns have long been thought to evolve under predator-driven selection, diversifying into phenotypes that reduce detection or warn predators of chemical defenses (Darwin 1859; Wallace 1889). For example, conspicuous wing coloration, often associated with prey defense, deters predators through learned avoidance of warning signals (aposematic cues that advertise a prey’s unpalatability or other defenses) (Ruxton et al. 2018). Conversely, wing shape modifications, such as hindwing tails, can increase survival by deflecting attacks away from vital body parts (Chotard et al. 2022). Despite the multifaceted selective pressures acting on antipredator defensive traits (Endler 1991; Kikuchi et al. 2023), macroevolutionary studies have seldom assessed the relative effects of wing shape and color pattern evolution simultaneously on the diversification of Lepidoptera (Puissant et al. 2026).

The evolution of aposematic coloration may promote speciation in butterflies (Mallet 1993), yet evidence remains limited to a few model systems (Kunte et al. 2021). In Ithomiini (clearwing butterflies), rates of mimicry gain and loss decline over time in species-rich clades, suggesting a slowdown in color-pattern turnover, consistent with ecological saturation (Chazot et al. 2025). However, speciation rates show no detectable association with these mimicry traits (Chazot et al. 2025). In contrast, the evolution of hindwing tails has been linked to rapid phenotypic diversification in *Papilio* swallowtails (Owens et al. 2020), and to increased diversification rates in non-diurnal moths (Rubin et al. 2018). A broader understanding of antipredator defense macroevolution can be gained by evaluating how multiple defensive traits jointly influence diversification across Lepidoptera lineages exhibiting extensive phenotypic variation.

To address the knowledge gaps of how different antipredator defenses influence diversification, we study the skipper butterfly subfamily Eudaminae (family Hesperiidae), a diverse clade of 595 recognized species (Warren et al. 2025) with an evolutionary history spanning over 35 million years (Espeland et al. 2018; Chazot et al. 2019; Kawahara et al. 2023). Eudaminae radiated across the Americas, primarily in the Neotropics, with several lineages occupying temperate regions, and a single genus, *Lobocla* Moore (10 recognized species) occurring in East and Southeast Asia (Toussaint et al. 2025). Eudaminae exhibit repeated evolution of similar traits, including dorsal blue-green iridescence and hindwing tails (Li et al. 2019), both potentially associated with antipredator defenses.

Blue-green coloration may advertise evasive flight behavior (Janzen et al. 2009), as suggested for other agile butterflies with bright blue markings like *Morpho* butterflies (Pinheiro and Freitas 2014; Pinheiro et al. 2016; Ledamoisel et al. 2025). Iridescence may also be involved in motion dazzling as an antipredator defense in fast-moving prey, where high-contrast color patterns blur predators’ perception of speed and trajectory (Stevens et al. 2008; Silvasti et al. 2024). Blue-green iridescence may also enhance camouflage (Thayer 1910), which can be enhanced by glossy backgrounds such as green vegetation in humid conditions (Kjernsmo et al. 2020; Thomas et al. 2023). Iridescence in Eudaminae likely represents a macroevolutionary trait optimum that correlates with increasing body size and morphometric proxies of increased flight speed (Linke et al. 2025a). Hindwing tails, in turn, can deflect attacks (Linke et al. 2025b), but may also trigger predator learning of traits associated with evasiveness (Linke et al. 2022) and/or unpalatability found in some tailed species of Eudaminae (Linke et al. 2024).

Eudaminae also show structured diel activity patterns (DeVries et al. 2008), with multiple lineages evolving strict crepuscular or nocturnal habits (Austin 2008). Environmental variation and exposure to distinct predator guilds may therefore have strongly influenced trait diversification in Eudaminae, as observed in other Lepidoptera (Nokelainen et al. 2024). We hypothesize that antipredator defenses, in interaction with environmental shifts (tropical *vs*. temperate occurrence; diurnal *vs*. non-diurnal activity), have influenced diversification in Eudaminae. Specifically, we predict that defensive traits (presence of hindwing tails and blue-green iridescent wing coloration) have evolved repeatedly through convergent evolution, shifted toward adaptive trait optima, and are associated with elevated speciation rates relative to lineages lacking these traits. We further predict that evolutionary losses of these traits may be more frequent than gains, reflecting trade-offs with flight performance, changes in predation pressure, or alternative defensive strategies. By integrating multiple traits and ecological factors, our study provides a novel mechanistic insight into how predation and environmental context shape phenotypic evolution and lineage diversification in butterflies.

## Materials and Methods

### Morphological, Ecological, and Geographic Data

We photographed Eudaminae specimens from natural history museums in Lima (Peru), Berlin (Germany), and London (United Kingdom), as well as at the Biology Centre CAS in České Budějovice (Czechia). We supplemented these photographs with online and published images of mounted butterflies with metric scales (Shuey 1991; Siewert et al. 2020; Zhang et al. 2023; Warren et al. 2025). In total, the dataset comprised 1,349 specimens representing 281 species (Table S1). Of those, 81% of the species had measurements from at least three specimens, whilst for the remainder, we could only measure a single or two specimens due to limited availability; the total median was five specimens per species (Table S1).

We quantified hindwing tail length relative to body size from three linear measures: 1) forewing length (distance between base and apex), 2) hindwing length (base to vein Cu_2_), and 3) hindwing tornus length (base to vein 2A) (Fig. 1). For tailed species represented by more than two males and females in our photographic dataset, no significant sexual dimorphism in hindwing tail ratio (tornus length divided by hindwing length) was detected (t-tests with adjusted p-values). As tail length relative to body size does not differ either significantly between males and females in the subtribe Eudamina (Linke et al. 2025a), measurements were averaged per species. Allometric effects were accounted for by extracting residuals from a log-log phylogenetic regression of hindwing tornus length against the forewing and hindwing lengths using the function “phyl.resid” in the R v. 4.4.1 (R Core Team 2024) package phytools (Revell 2024). These residuals represented phylogenetically corrected tail length per species. In addition, we scored the presence of hindwing tails and blue-green wing coloration for all described Eudaminae species, including those not in our photographic database, using an online photographic catalogue (Warren et al. 2025) and species descriptions (Bertrand et al. 2014; Siewert et al. 2015, 2018a, 2018b). Blue-green coloration was considered present when clearly visible on the dorsal wings, whereas ambiguous or negligible blue-green coloration only on the thorax was scored as absent.

**Figure 1:**
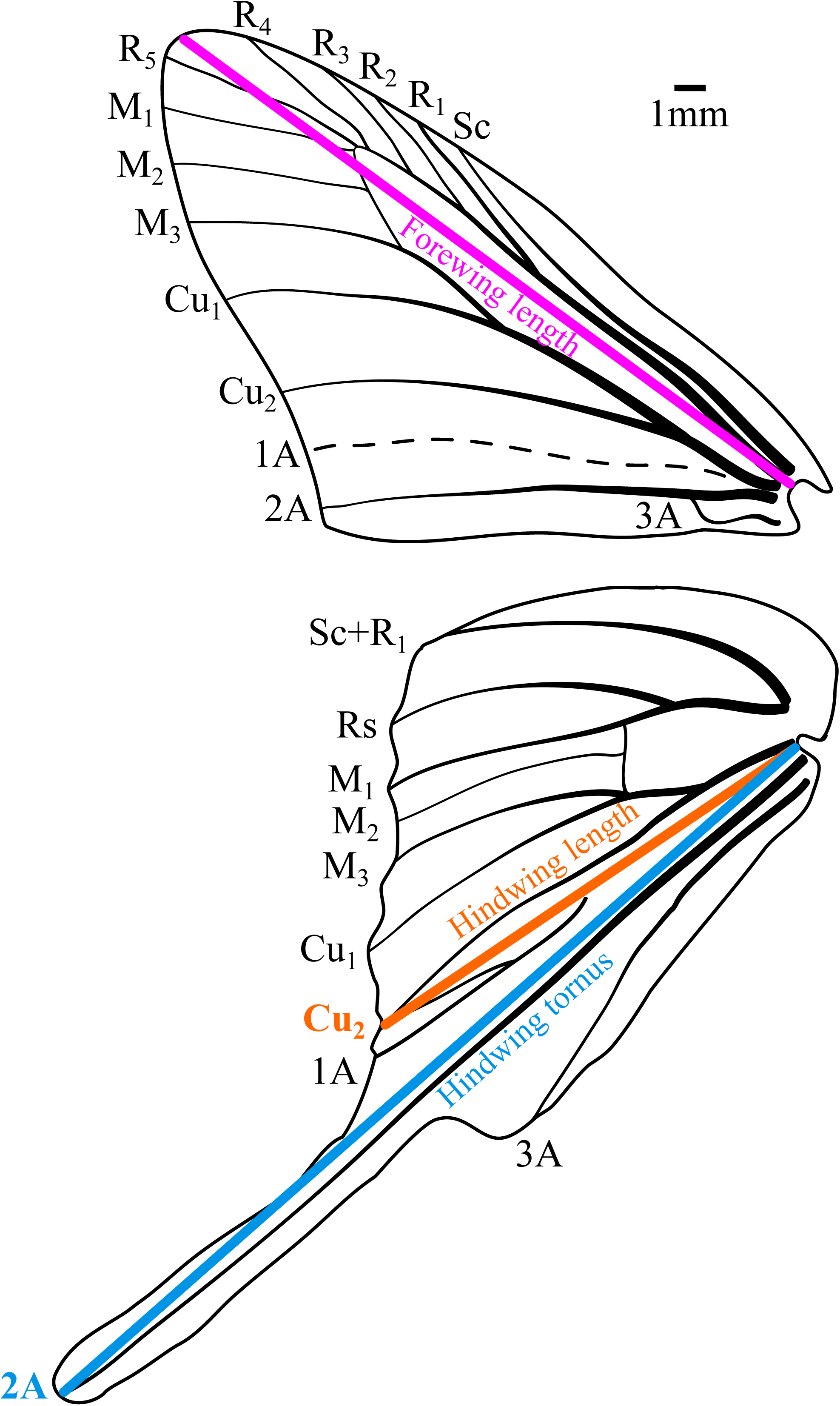
The linear measures of forewing and hindwings consisted of 1) forewing length, distance between the wing base and apex, 2) hindwing length, distance between the wing base and vein Cu_2_, and 3) hindwing tornus, distance between the base and vein 2A. Wing venation of *Urbanus proteus*.

We compiled diel activity information from published records (Mielke 1967; Austin and Mielke 2000, 2008; Burns and Janzen 2005; Austin 2008; DeVries et al. 2008; Sourakov and Houlihan 2017; Núñez Bustos 2018) and classified species as diurnal or non-diurnal (crepuscular/nocturnal flight). For species with limited diel activity data, e.g., those occasionally collected in light traps (Burns et al. 2013) or roosting in caves (Viloria 1993), we used the presence of red-eye pigmentation as a proxy for non-diurnal activity (Toussaint and Warren 2019).

We determined the distribution of Eudaminae species outside tropical bioregions defined in Morrone (2014), using an online database of range maps (Savela 2025), textual regional information (Warren et al. 2025), metadata gathered from museum collections, and species descriptions (Pfeiler et al. 2018). Geo-referenced occurrences (GBIF 2024) cleaned with CoordinateCleaner (Zizka et al. 2019) and Research Grade observations in iNaturalist (https://www.inaturalist.org/) were used to confirm the ranges of extratropical species. Species occurring outside the tropics were classified as either widespread (occasional or seasonal strays out of the tropics) or established breeders (with confirmed oviposition or immature stages outside the tropics). These categories were assigned by consulting online nature guides, life history databases (Cary and Toliver 2024; Lotts et al. 2024; NatureServe 2024), and species descriptions.

All morphological, ecological, and geographic information was consolidated into a comprehensive species-level trait database of Eudaminae including: 1) hindwing tail presence/absence; 2) presence/absence of blue-green wing coloration; 3) diel activity (diurnal/non-diurnal); and 4) geographic category (tropical *vs*. non-tropical established breeder or widespread) (Table S2).

### Taxon Sampling and DNA Sequencing

We sampled 184 skipper butterflies between 1992 and 2021 from localities across North, Central and South America (Table S3). Genomic DNA was extracted from two to three legs or abdomens using DNeasy Blood & Tissue kits (QIAGEN, Hilden, Germany). For 101 individuals, we enriched 425 loci using the BUTTERFLY probe kit 1.0 (Espeland et al. 2018), and for two individuals, we enriched 13 protein-coding genes using the BUTTERFLY probe kit 2.0 (Kawahara et al. 2018). Target enrichment, library preparation, and 150 bp paired-end read sequencing on Illumina NovaSeq platforms were carried out by RapidGenomics (Florida, USA). We also generated whole-genome sequencing data for 81 individuals, targeting ∼7 Gb raw data per sample (Hesperiidae genome sizes: ∼293–645 Mb (Liu et al. 2020). Library preparation and 150 bp paired-end read sequencing on Illumina NovaSeq6000 platforms were conducted by Novogene Europe (Cambridge, UK).

We complemented our dataset with published whole-genome sequences (Li et al. 2019; Ribeiro et al. 2021, 2025) and target-capture alignments based on the BUTTERFLY probe kits 1.0 and 2.0 (Kawahara et al. 2023; Toussaint et al. 2025). Overall, our taxon sampling included 281 Eudaminae species and 29 outgroup taxa from other Hesperiidae subfamilies.

### Data assembly and DNA alignments

Raw Illumina reads from target capture and whole-genome sequencing were processed in SECAPR v. 2.2.3 (Andermann et al. 2018; Ribeiro et al. 2021). Adapters and bases with Phred scores < 20 were removed using Trimmomatic v. 0.39 (Bolger et al. 2014), and read quality was assessed with FastQC v. 0.11.9 (Andrews 2010). *De novo* contigs were assembled with SPAdes v. 3.14.1 (Bankevich et al. 2012) using k-mer sizes of 21, 33, 55 and 77.

We retrieved 425 loci of interest (Espeland et al. 2018) from *de novo* contigs using the SECAPR ‘find_target_contigs’ function, which uses LASTZ v. 1.04 (Harris 2007). Loci were aligned with MAFFT v. 7.130 (Katoh and Standley 2013) and trimmed to exon boundaries. Published alignments from (Kawahara et al. 2023) were incorporated using the MAFFT options ‘--addfull’ and ‘--keeplength’.

### Phylogenetic Analyses

We selected 369 of the 425 targeted loci, including 14 legacy genes widely used in butterfly systematics (Wahlberg and Wheat 2008; Kawahara et al. 2018), and loci with > 70% species coverage. The concatenated dataset was partitioned by gene and codon position, and PartitionFinder v. 2.1.1 (Lanfear et al. 2017) was used to estimate the optimal scheme under the GTR+Γ model using Bayesian Information Criterion (BIC) and the ‘rcluster-percent’ search algorithm set to 10%.

Maximum-likelihood phylogenies were inferred in IQ-TREE v. 2.2.2.6 (Hoang et al. 2018a; Minh et al. 2020) using 100 parsimony starting trees and five independent runs. Branch support was assessed with 1,000 ultrafast bootstrap replicates (UFBoot; Hoang et al. 2018a), each optimized by NNI, and 1,000 Shimodaira-Hasegawa-like approximate likelihood-ratio test replicates (SH-aLRT; Guindon et al. 2010). Substitution models for each partition were selected with IQ-TREE’s ModelFinder (Kalyaanamoorthy et al. 2017) using BIC. Bayesian phylogenetic inference was conducted in ExaBayes v. 1.5.1 (Aberer et al. 2014) using GTR+Γ models for all partitions. Two independent runs of one million generations were performed, each with one cold and three heated chains and sampling every 100 generations. Convergence was assessed in ExaBayes based on the average standard deviation of split frequencies, with values below 1% considered indicative of convergence. After discarding the first 25% of sampled trees as burnin, we estimated an extended majority-rule consensus tree.

We inferred a species tree under the summary multispecies coalescent approach using weighted ASTRAL (wASTRAL; Zhang and Mirarab 2022) as implemented in ASTER v. 1.23 (Zhang et al. 2025). Because accurate gene trees are essential for summary-coalescent methods and maximum parsimony performs well under incomplete lineage sorting (Vanderpool et al. 2020), we inferred parsimony gene trees for all loci. Taxa with > 50% missing data per locus were removed using Sequence_Cleaner (https://github.com/metageni/Sequence-Cleaner). Gene trees were inferred in MPBoot v. 1.1 (Hoang et al. 2018b) with 1,000 UFBoot replicates, and nodes with < 50% support were collapsed using Newick Utilities v. 1.6 (Junier and Zdobnov 2010). The wASTRAL species tree was then estimated from the vetted gene trees, with branch supports estimated using local posterior probabilities.

To quantify topological congruence among phylogenetic inferences, we calculated quartet distances using the R package Quartet (Smith 2024). For this analysis, poorly supported nodes were collapsed using thresholds of < 95% UFBoot and SH-aLRT (IQ-TREE), < 0.95 posterior probability (ExaBayes), and < 0.5 local posterior probability (wASTRAL).

### Divergence Time Estimation

Because tree topologies were highly congruent across phylogenetic approaches, we time-calibrated the bifurcating IQ-TREE phylogeny using eight secondary calibrations consistent with fossil-based butterfly chronograms (Espeland et al. 2018; Chazot et al. 2019; Kawahara et al. 2023). Calibration bounds followed the 95% highest posterior density (HPD) reported in (Kawahara et al. 2023), and we constrained the crown ages of: 1) Hesperiidae (68–77 Myr), 2) Tagiadinae (43–51 Myr), 3) Pyrginae (39–49 Myr), 4) Hesperiinae (37–47 Myr), 5) Hesperiini (25–34 Myr), 6) Eudaminae (32–41 Myr), 7) the most recent common ancestor (MRCA) of Hesperiinae + Trapezitinae (43–52 Myr), and 8) the MRCA of Heteropterinae + Hesperiinae + Trapezitinae (46–55 Myr).

Divergence times were estimated under the Bayesian independent-rates relaxed clock model (Rannala and Yang 2007) in MCMCTree, part of PAML v. 4.10.5 (Yang 2007). First, we used BASEML (in PAML) to estimate an informed prior on the mean substitution rate for the 369-loci concatenated alignment, represented by a gamma-Dirichlet distribution with α = 2 and β = 76.8. Second, we calculated the gradient and Hessian matrix of branch lengths using the approximate likelihood calculation method (dos Reis and Yang 2011). Third, we ran MCMCTree under the available HKY85+Γ model and with uniform time priors for all eight calibrations. Four independent MCMC runs of 1 million generations were performed, sampling every 100 generations after a burnin of 10,000 generations. Convergence and Effective Sample Size (ESS) > 200 were confirmed using Tracer v. 1.7.2 (Rambaut et al. 2018).

We also jointly inferred topology and divergence times using BEAST v. 2.7.7 (Bouckaert et al. 2014). Due to computational constraints, this analysis included 84 loci: 14 legacy genes and 70 loci with < 50% missing data and > 75% species coverage. Sequences were concatenated and partitioned into two subsets: first/second codon positions and third codon positions. We applied the Optimized Relaxed Clock model (Douglas et al. 2021) with a Gamma prior distribution on the mean clock rate (mean = 0.001, β = 1.0) and estimated site models using bModelTest v. 1.3.3 (Bouckaert and Drummond 2017). A Birth-Death tree model and the same eight calibration constraints, normally distributed around the 95% HPD intervals of Kawahara et al. (2023), were used to infer absolute time. Four MCMC runs of 80–100 million generations were performed, sampling every 20,000 generations. After discarding non-stationarity states, the combined posterior samples showed ESS values > 200, and we summarized the trees into a maximum clade credibility (MCC) tree. Finally, we repeated the BEAST analysis using an alternative set of 70 loci ranked by clock-likeness and phylogenetic information content using SortaDate (Smith et al. 2018) (Fig. S1).

### Ancestral Range Estimation, Frequency and Directionality of Dispersal

We classified the extant distribution of Eudaminae species into seven bioregions: 1) Mesoamerica and the Pacific slope of the Andes, 2) the Antilles, 3) the Andes above ∼1,000 m, 4) Amazonia and the eastern foothills of the Andes, and 5) the Atlantic Forest, as well as two extratropical regions, 6) the Nearctic and 7) the Pampas (Table S4). The five tropical regions roughly correspond to those of Morrone (2014). Because the Asian genus *Lobocla* represents a single dispersal event out of the Americas (Li et al. 2019; Toussaint et al. 2025), it was excluded from biogeographic analyses.

We inferred ancestral ranges using the Dispersal-Extinction-Cladogenesis (DEC) model (Ree and Smith 2008) implemented in the R package BioGeoBEARS (Matzke 2013). To avoid unrealistic distributions, ranges spanning highly disjunct bioregions were excluded. Specifically, we disallowed range combinations that included the Nearctic, Mesoamerica, and/or the Antilles together with the Atlantic Forest and/or Pampas, without including Amazonia or the Andes, which represented the primary areas connecting northern and southern Neotropical regions. We estimated dispersal dynamics with Biogeographical Stochastic Mapping (BSM; Dupin et al. 2017), performing 100 stochastic mappings on each of 50 randomly selected posterior trees from the MCMCTree and BEAST analyses (5,000 BSM replicates per dating framework). We recorded the frequency and direction of dispersal events by averaging BSM replicates, and summarized them across one-million-year time bins using scripts from Matos-Maraví et al. (2021).

### Testing Adaptive Evolution of Antipredator Traits Across Environmental Regimes

We tested whether environmental variation and diel activity influenced hindwing tail evolution. Species were assigned to three environmental regimes based on the consolidated Eudaminae trait database (Table S2): 1) tropical diurnal, 2) tropical non-diurnal, and 3) temperate diurnal (all temperate breeders are diurnal). We fitted Ornstein-Uhlenbeck (OU) models to examine whether the phylogenetically size-corrected hindwing tail lengths evolved toward selective optima (θ) under these regimes (Hansen 1997; Butler and King 2004), estimating the rate of stochastic evolution (σ^2^) and the strength of selection (α) toward the optima (Beaulieu et al. 2012). Six evolutionary models were compared: 1) OU1, a single θ and shared σ^2^ and α across selective regimes; 2) OUM, distinct θ per regime; 3) OUMV, distinct θ and σ^2^; 4) OUMA, distinct θ and α; 5) BM, non-adaptive Brownian motion with a single σ^2^; and 6) BMS, Brownian motion with distinct σ^2^ per regime.

Analyses were performed on both MCMCTree and BEAST time-calibrated phylogenies using the R package OUwie (Beaulieu et al. 2012). Environmental regimes were reconstructed with 100 stochastic character mappings using the root prior method of FitzJohn et al. (2009) and the All-Rates-Different model in phytools (Revell 2024). Model support was assessed using Akaike weights averaged across all stochastic mappings.

To evaluate the effect of blue-green iridescent coloration on hindwing tail evolution, we repeated OU analyses using the presence/absence of blue-green dorsal coloration as the selective regime. This allowed us to test how an additional antipredator trait may influence tail evolution under environmental variation. In addition, to count the number of hindwing tail and blue-green coloration gains and losses, we reconstructed 100 stochastic character mappings on these traits separately using the All-Rates-Different model in phytools (Revell 2024).

### Testing Trait-Dependent Diversification Associated with Antipredator Defenses

We evaluated whether hindwing tails or blue-green coloration, in combination with environmental regimes, influenced diversification rates using the Hidden State-dependent Speciation and Extinction (HiSSE) framework (Beaulieu and O’Meara 2016). Because hindwing tail length is continuous, we discretized it using the residuals of the log-log phylogenetic regression of tail length against wing size (range: 0.134 to 0.689; Table S4) and applied a cutoff value of 0.25. Based on environmental and phenotypic attributes, we defined four discrete character states: 1) TDO, tropical diurnal species lacking the focal trait (hindwing tails or blue-green coloration); 2) TNO, tropical non-diurnal species lacking the trait; 3) TDW, tropical diurnal species expressing the trait; and 4) CDO, temperate diurnal species lacking the trait. We expanded the HiSSE model with two hidden character states. To reduce character space and improve computational feasibility, we excluded temperate breeders with hindwing tails (3 species) or blue-green coloration (2 species), as well as non-diurnal species with blue-green coloration (2 species). Widespread species (occasional strays into temperate regions; 23 species) were considered tropical, with supplementary analyses also run after excluding them to evaluate sensitivity to range assignments.

We fitted two HiSSE models in RevBayes v. 1.2.4 (Höhna et al. 2016), running two independent MCMC chains of 10,000 generations with tuning every 100 generations. After burnin and merging chains, all parameters showed ESS values > 200. Sampling probabilities were specified to account for incomplete taxon sampling, though trait-specific sampling differences could not be implemented. Analyses were repeated on both time-calibrated phylogenies (MCMCTree and BEAST). During the MCMC, ancestral states and stochastic character mappings were generated to quantify transition patterns among the four phenotype-environment categories. To assess differences in speciation rates among states, we computed pairwise posterior probabilities by calculating the proportion of posterior estimates in which one state’s speciation rate exceeded another. Posterior distributions of pairwise rate differences were summarized with 95% credible intervals (CI), with ranges excluding zero indicating strong differences in speciation rates between states.

### Testing Asymmetry in Evolutionary Gains and Losses of Antipredator Traits

We examined the frequency and directionality of transitions among the four phenotype-environment states used in the HiSSE analyses (TDO, TNO, TDW, CDO) by fitting alternative models of discrete trait evolution. Seven competing models were evaluated: 1) full model, all transition rates free to vary; 2) symmetric model, forward and backward transitions constrained to be equal; 3) equal-rates model, all transition rates equal; 4) no simultaneous shifts in wing phenotype and diel activity (no transitions between TDW and TNO); 5) no simultaneous shifts in wing phenotype, diel activity and geographic region (no transitions between TDW and TNO or between TDW and CDO); 6) no simultaneous shifts in diel activity, wing phenotype and geographic region (no transitions between TNO and TDW or between TNO and CDO); and 7) transitions restricted to the tropical diurnal state lacking the antipredator phenotype (TDO) (Fig. S2).

Transition models were fitted to both dated phylogenies (MCMCTree and BEAST) using the fitMK function in phytools (Revell 2024). We used the root prior method of FitzJohn et al. (2009) and accounted for divergence time and phylogenetic uncertainty by running each model across 100 posterior trees. Model support was evaluated using Akaike weights averaged across all trees.

## Results

### Morphological, Ecological, and Geographic Data

Among the 595 recognized species in Eudaminae, we identified 104 with hindwing tails and 164 with blue-green wing coloration. We documented non-diurnal activity (crepuscular/nocturnal) for 88 species and recorded 82 extratropical species, of which 48 are established breeders outside of the tropics. Non-diurnal Eudaminae have tailless hindwings and only the crepuscular genus *Phareas* Westwood (2 species) presents iridescent dorsal wings (Toussaint and Warren 2019). Trait representation in the time-calibrated phylogenies of 281 species was proportional, with 55 species exhibiting hindwing tails, 88 blue-green coloration, 45 non-diurnal activity, and 49 extratropical, including 26 temperate breeders (Table S4).

### Phylogenetic Analyses

The final concatenated DNA sequence alignment comprised 203,277 bp across 369 loci, with < 30% missing data per locus and 85,296 parsimony-informative sites. Phylogenies inferred with maximum likelihood (IQ-TREE), Bayesian inference (ExaBayes) and summary coalescence (wASTRAL) were highly congruent (Figs S3–S5). Node support was uniformly strong: 86% of nodes had > 95% UFBoot and SH-aLRT support, 93% had > 0.95 posterior probability in ExaBayes, and 92% had > 0.5 local posterior probability in wASTRAL. Quartet comparisons showed minimal topological conflict among the IQ-TREE, ExaBayes, and wASTRAL trees, with proportions of non-conflicting nodes and explicitly agreeing nodes close to one, and quartet divergence of ExaBayes and wASTRAL from the IQ-TREE topology close to zero.

Most genera were monophyletic except *Telemiades* Hübner, which was polyphyletic. *Telemiades delalande* (Latreille) consistently grouped with *Polygonus* Hübner, while *T. litanicus* (Hewitson) was embedded within *Ectomis* Mabille. To ensure generic monophyly, we here revise species classifications as follows:

*Polygonus delalande* (Latreille, [1824]), comb. n.; former generic placement, *Telemiades*

Hübner, [1819].

Synonyms:

*Telemiades delalande* (Latreille, [1924]).

*Hesperia delalande* Latreille, [1924].

*Pterygospidea panthea* Hewitson, 1868.

*Achlyodes amaurus* Mabille, 1889.

*Echelatus lucina* Schaus, 1913, p. 358, pl. 54, fig. 8.

***Ectomis litanicus* (Hewitson, 1876), comb. n.**; former generic placement, *Telemiades* Hübner, [1819].

Synonyms:

*Telemiades litanicus* (Hewitson, 1876).

*Eudamus litanicus* Hewitson, 1876, p. 354.

### Divergence Time Estimation

BEAST trees were highly congruent with the IQ-TREE topology (explicitly agreeing nodes = 0.99; quartet divergence = 0.002) (Figs S6–S7). The supplementary BEAST analysis using 70 clock-like loci and 14 legacy genes resulted in nearly identical trees (Fig. S1), although two species were not represented in this dataset. Divergence time estimates were highly consistent across approaches (Table 1) and, as expected, aligned with the 95% HPD intervals of recent butterfly chronograms (Chazot et al. 2019; Kawahara et al. 2023) (Fig. 2). However, our estimates were slightly older than those reported by Toussaint et al. (2025), likely because the latter study used younger secondary calibrations derived from Kawahara et al. (2019).

**Figure 2:**
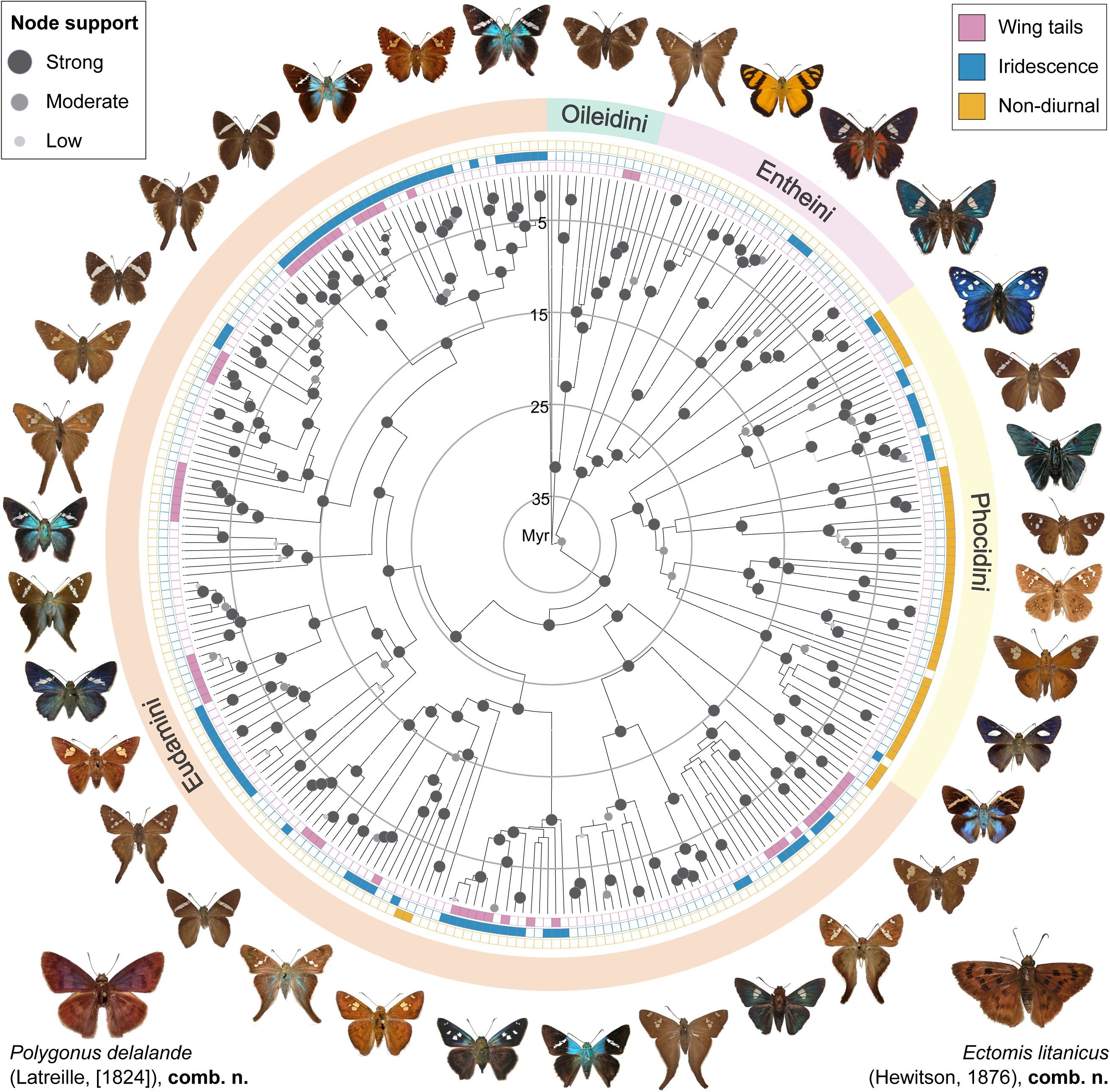
Time-calibrated tree of Eudaminae inferred with BEAST (scale in million of years, Myr). A total of 84 loci were used. Node support evaluated across four phylogenetic inference methods: IQ-TREE, ExaBayes, wASTRAL and BEAST. A node is considered strongly supported within each method when it showed > 95% UFBoot and SH-aLRT (IQ-TREE), > 0.95 posterior probabilities (ExaBayes and BEAST), and > 0.5 local posterior probability (wASTRAL). These support metrics are summarized into three categories mapped on the tree: strong (supported by all four methods), moderate (supported by three methods), and low (supported by two methods). Wing traits and flight activity (hindwing tails, blue-green coloration, and non-diurnal behavior) mapped onto the phylogeny to illustrate convergent evolutionary patterns. Butterfly images from (Warren et al. 2025) are arranged according to taxonomic tribe and phylogenetic relationships. In the light of phylogenomic data, the taxonomic status of *Polygonus delalande* and *Ectomis litanicus* (Eudamini, Telemiadina) are revised.

**Table 1:** Divergence time estimates for Eudaminae tribes. Crown ages and 95% highest posterior density intervals (in million years) are summarized from previously published butterfly trees and from our estimates using BEAST and MCMCTree. BEAST results are shown for two datasets: (i) the main analysis maximizing sequence completeness, and (ii) an alternative analysis using SortaDate-selected clock-like loci.

| *Clade* | *Chazot et al. 2019* | *Kawahara et al. 2023* | *Toussaint et al. 2025* | *BEAST* | *BEAST, alternative* | *MCMCTree* |
| --- | --- | --- | --- | --- | --- | --- |
| *Eudaminae* | 35.2 [28.8–42.6] | 35.2 [32.4–40.9] | 33.9 [32.1–35.9] | 40.2 [37.7–42.9] | 40.3 [37.9–42.9] | 41.0 [40.5–42.6] |
| *Oileidini* | - | 24.7 [21.4–31.3] | 29.2 [25.1–32.5] | 31.8 [28.0–35.6] | 32.6 [28.8–36.2] | 34.6 [31.6–37.4] |
| *Entheini* | 28.3 [23.0–35.0] | 26.4 [24.2–31.9] | 26.1 [24.6–27.8] | 31.7 [29.0–34.5] | 30.7 [28.0–33.6] | 33.3 [31.4–35.3] |
| *Phocidini* | 26.3 [21.5–32.0] | 26.2 [24.5–32.4] | 25.0 [23.8–26.4] | 30.0 [27.8–32.4] | 29.4 [26.9–31.6] | 33.1 [31.0–34.8] |
| *Eudamini* | 27.4 [22.5–33.3] | 26.1 [24.6–30.7] | 26.6 [25.3–28.0] | 31.4 [29.2–33.7] | 31.5 [29.2–33.7] | 33.9 [32.0–35.4] |

### Ancestral Range Estimation and Frequency and Directionality of Dispersal

Biogeographical inferences indicated an Amazonian origin and subsequent diversification for early diverging Eudaminae lineages, followed by repeated dispersal into Mesoamerica, the Andes, and the Atlantic Forest (Fig. S8). Dispersal into extratropical regions increased sharply from the mid-Miocene onward. Pampas lineages originated mainly from the Atlantic Forest and Amazonia during the last 5 Myr. Nearctic lineages arose primarily from Mesoamerica, and to a lesser extent from the Antilles, during the last 15 Myr (Fig. S9). Dispersal from extratropical back into tropical regions also increased during the mid-Miocene, but at substantially lower rates than dispersal into the Nearctic and Pampas (Fig. 3).

**Figure 3:**
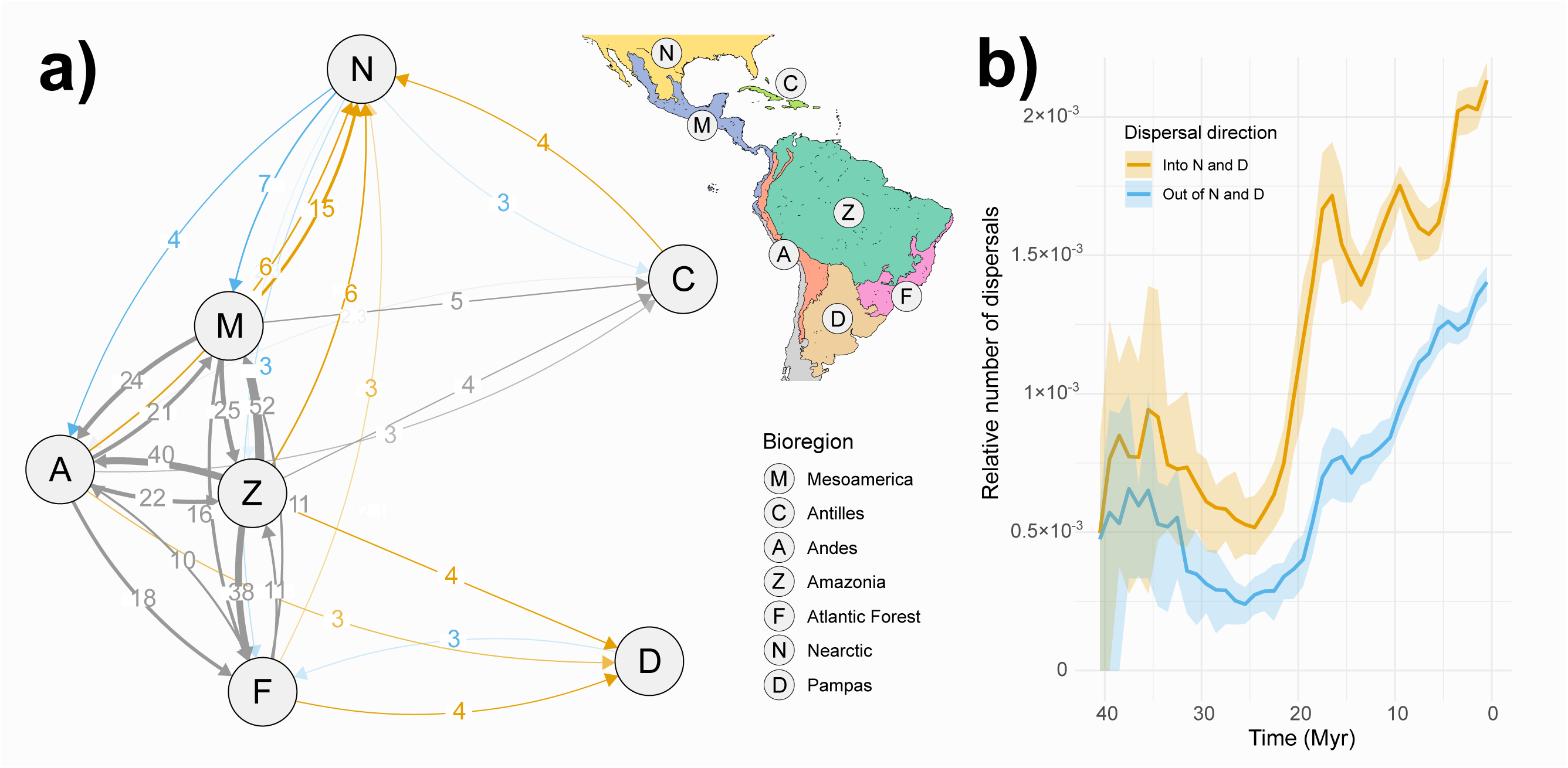
Dispersal dynamics of Eudaminae in the Americas estimated using the time-calibrated BEAST phylogeny and Biogeographical Stochastic Mapping (BSM) with BioGeoBEARS. a) Estimated dispersal events among biogeographical regions. Arrows indicate the direction and number of inferred dispersal events. Only connections with more than two events are shown. Node layout reflects the strength of biogeographical interconnections, arranged using the force-embedded algorithm in the R package qgraph (Epskamp et al. 2012). Dispersal events into and out of extratropical regions (Nearctic and Pampas) are highlighted in orange and blue, respectively. b) Temporal patterns of dispersal into and out of extratropical regions over millions of years (Myr). The number of dispersal events is normalized by lineage counts within each time bin (1 Myr). Mean values and the 25%–75% quantiles are derived from 5,000 BSM replicates, accounting for phylogenetic and biogeographical uncertainty.

### Adaptive Evolution of Hindwing Tails Across Environmental Regimes, but not of Blue-Green Coloration

Hindwing tails in Eudaminae evolved at least seven times and blue-green coloration, fifteen times. To test the adaptive versus neutral evolution of hindwing tails under environmental variation, we compared six OU and BM models. The OUMV model, allowing distinct trait optima (θ) and evolutionary rates (σ^2^) among regimes, was the best-supported model for both the MCMCTree (average Akaike weight = 0.70, standard error (SE) = 0.05) and BEAST phylogenies (average Akaike weight = 0.75, SE = 0.04). This model indicated significantly longer hindwing tails in tropical diurnal species relative to both tropical non-diurnal and temperate diurnal species (Table S5), consistent with divergent selective optima among environments.

In contrast, hindwing tail evolution did not show strong selective differentiation in relation to blue-green wing coloration. The BMS model was best supported (average Akaike weights: 0.60 for MCMCTree; 0.52 for BEAST), indicating neutral rather than adaptive trait divergence across coloration regimes (Table S6).

### Elevated Speciation in Diurnal Lineages with Hindwing Tails and Blue-Green Coloration

HiSSE analyses revealed that tropical non-diurnal species had the lowest speciation rates, whereas tropical diurnal species with hindwing tails exhibited the highest rates among the examined states (Fig. 4a). Pairwise comparisons indicated that non-diurnal species had lower speciation rates than tropical diurnal species (posterior probabilities > 0.95). The strongest contrast was between tailed tropical diurnal species and non-diurnal species, with the 95% CI excluding zero, while non-tailed tropical diurnal species likely had higher rates than non-diurnal species, but their 95% CI marginally overlapped zero only with the BEAST tree (range: 0.001 to 0.086 for MCMCTree; 0.003 to 0.089 for BEAST). Evidence for higher speciation in tailed *vs*. non-tailed tropical diurnal species was moderate (posterior probabilities = 0.83 for MCMCTree, 0.86 for BEAST), but the 95% CI included zero (Table 2). The macroevolutionary dynamics remain virtually the same when excluding widespread species (i.e., species with occasional or seasonal strays out of the tropics) from the analyses (Table S7).

**Figure 4:**
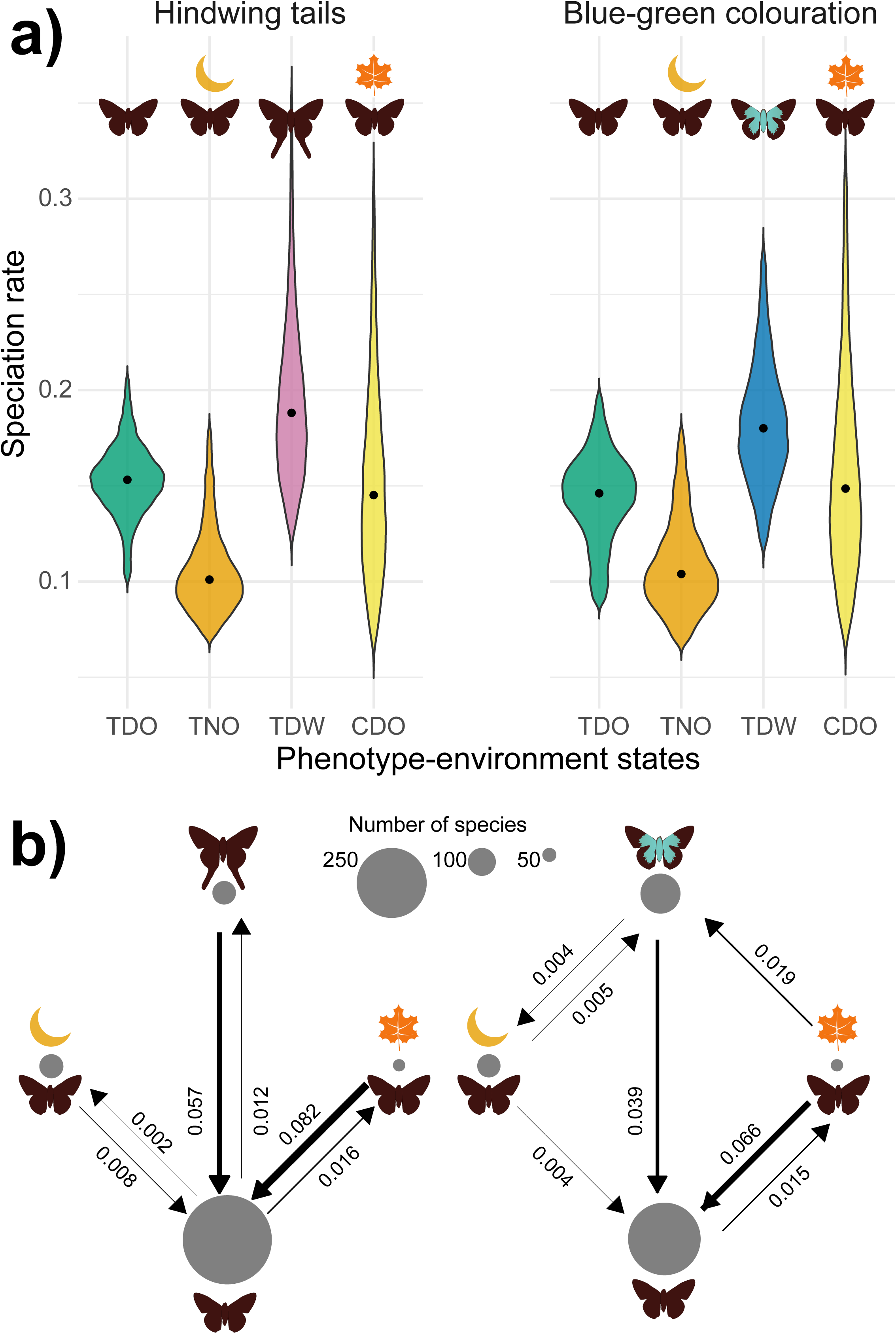
Speciation rates estimated using the Hidden State-dependent Speciation and Extinction (HiSSE) framework in RevBayes, and transition rates using the fitMK function in phytools. Results are shown for the BEAST tree; analyses with MCMCTree produced similar patterns (Tables S5–S7). Four discrete states combine environmental and phenotypic attributes: TDO, tropical diurnal species lacking the focal trait (hindwing tails or blue-green coloration); TNO, tropical non-diurnal species lacking the trait; TDW, tropical diurnal species expressing the trait; CDO, temperate diurnal species lacking the trait. a) Posterior distributions (95% HPD) of speciation rates for each state estimated using HiSSE. b) Transition rates between states under the best-fit models: restricted transitions for hindwing tails (only involving the tailless tropical diurnal state) and all-rates-different for blue-green coloration. The number of species in each state is indicated as proportional circles.

**Table 2:** Speciation rate estimates under the Hidden State-dependent Speciation and Extinction (HiSSE) framework implemented in RevBayes. Four discrete states combine environmental and phenotypic attributes: TDO, tropical diurnal species lacking the focal trait (hindwing tails or blue-green coloration); TNO, tropical non-diurnal species lacking the trait; TDW, tropical diurnal species expressing the trait; CDO, temperate diurnal species lacking the trait. Pairwise differences in speciation rates (λ) summarized with 95% credible intervals. Comparisons whose intervals do not overlap zero are marked with an asterisk. Differences are reported as state_1_ state_2_, so positive values indicate higher speciation rates in the state 1. Posterior probabilities represent the proportion of posterior samples in which the speciation rate of one state exceeded that of the other.

| *Comparison* | *Rate difference*<br>*(MCMCTree)* | *Posterior probability*<br>*(MCMCTree)* | *Rate difference*<br>*(BEAST2 tree)* | *Posterior probability*<br>*(BEAST2 tree)* |
| --- | --- | --- | --- | --- |
| *Hindwing tails* |  |  |  |  |
| $\lambda_{TDW} \square \lambda_{TNO}$ | 0.076 [0.020–0.168]* | 0.993 | 0.083 [0.019–0.228]* | 0.993 |
| $\lambda_{TDO} \square \lambda_{TNO}$ | 0.049 [0.001–0.086]* | 0.977 | 0.048 [□0.003–0.089] | 0.969 |
| $\lambda_{TDW} \square \lambda_{TDO}$ | 0.029 [□0.025–0.120] | 0.835 | 0.036 [□0.024–0.178] | 0.863 |
| $\lambda_{CDO} \square \lambda_{TNO}$ | 0.038 [□0.042–0.192] | 0.808 | 0.041 [□0.052–0.192] | 0.804 |
| $\lambda_{TDW} \square \lambda_{CDO}$ | 0.038 [□0.114–0.152] | 0.747 | 0.044 [□0.106–0.198] | 0.752 |
| $\lambda_{TDO} \square \lambda_{CDO}$ | 0.010 [□0.141–0.087] | 0.576 | 0.006 [□0.143–0.090] | 0.546 |
| **Blue-green coloration** |  |  |  |  |
| $\lambda_{TDW} \square \lambda_{TNO}$ | 0.072 [0.024–0.145]* | 0.995 | 0.073 [0.020–0.141]* | 0.994 |
| $\lambda_{TDO} \square \lambda_{TNO}$ | 0.037 [□0.008–0.082] | 0.949 | 0.035 [□0.015–0.082] | 0.933 |
| $\lambda_{TDW} \square \lambda_{TDO}$ | 0.036 [□0.015–0.108] | 0.920 | 0.038 [□0.013–0.105] | 0.928 |
| $\lambda_{CDO} \square \lambda_{TNO}$ | 0.039 [□0.042–0.186] | 0.829 | 0.042 [□0.052–0.206] | 0.813 |
| $\lambda_{TDW} \square \lambda_{CDO}$ | 0.032 [□0.107–0.134] | 0.738 | 0.031 [□0.130–0.134] | 0.704 |
| $\lambda_{CDO} \square \lambda_{TDO}$ | 0.002 [□0.078–0.146] | 0.519 | 0.006 [□0.080–0.169] | 0.551 |

For blue-green wing coloration, evidence for state-dependent diversification was weaker than found for hindwing tail evolution (Fig. 4a). Diurnal species with blue-green wings had higher speciation rates than non-diurnal species (posterior probability = 0.99; 95% CI excluded zero in both MCMCTree and BEAST trees). Pairwise comparisons indicated moderate support for higher speciation rates in blue-green wing coloration lineages than diurnal species lacking such a wing color pattern (posterior probabilities = 0.90–0.95, but 95% CI crossed zero) (Table 2).

### Frequent Losses Indicate Evolutionary Lability of Hindwing Tails and Blue-Green Coloration

The best-fit transition model for hindwing tail evolution restricted transitions to those involving the non-tailed tropical diurnal state (TDO), supported by both MCMCtree and BEAST phylogenies (mean Akaike weight ∼0.45). Estimated rates were highest for shifts from temperate diurnal to tropical diurnal states, and for loss of hindwing tails in tropical diurnal species.

Reverse transitions were lower but still detectable, involving the evolution of hindwing tails in tropical diurnal species and shifts to extratropical regions. The model also suggested that non-diurnal lineages evolved from tropical, non-tailed ancestors (Fig. 4b).

The best-fit model was all-rates-different for blue-green wing coloration evolution (mean Akaike weights ∼0.55 across dated phylogenies). Transition rates for loss of blue-green coloration were higher than its evolution, which was primarily from temperate diurnal species and from non-diurnal tropical species. Unexpectedly, transition rates from tropical diurnal species lacking the trait toward the evolution of blue-green coloration were negligible. The highest transition rates were for shifts from temperate to tropical regions. Non-diurnal species evolved toward diurnality at higher rates than the reverse (Fig. 4b).

## Discussion

Using phylogenomic, morphometric, and comparative methods, we tested the hypothesis that antipredator defenses, in interaction with environmental variation, promoted species diversification in Eudaminae. Our results largely corroborate this hypothesis. As predicted, defensive traits showed repeated convergent evolution, shifts toward adaptive trait optima, and associations with increased diversification rates. Convergent evolution of hindwing tails and blue-green coloration on wings is widespread and partly driven by selection, particularly for hindwing tails, which evolve toward distinct trait optima rather than through neutral drift. Both hindwing tails and blue-green wing coloration were associated with elevated speciation rates relative to non-diurnal lineages lacking these traits. However, evolutionary transitions away from these traits were more common than their acquisition. This pattern indicates that, while antipredator defenses may enhance diversification, they are also evolutionarily labile.

### Environmental Context Shapes Adaptive Gains and Frequent Losses of Antipredator Defenses

The lability of conspicuous antipredator phenotypes likely reflects ecological trade-offs or fluctuating predator-driven selection. For example, patterns of repeated gain and loss of wings in stick insects (order Phasmatodea) suggest that even complex insect traits can be evolutionarily flexible and highly responsive to changing ecological pressures, rather than permanently stable features (Forni et al. 2022). High transition rates toward loss of hindwing tails and blue-green coloration in tropical diurnal lineages suggest that phenotypes lacking these traits may represent macroevolutionary “sinks” (Loeffler-Henry et al. 2023). Eudaminae encounter diverse avian predators, including aerial hunters such as kingbirds, tyrant flycatchers, and vireos, as well as opportunistic predators such as tanagers (Pinheiro 1996; Linke et al. 2025b). Hindwing tails may incur aerodynamic costs or vary in their efficacy in deflecting attacks depending on prey body size (Rubin et al. 2018) and predator ability to catch evasive prey (Linke et al. 2025b). These context-dependent selective pressures may explain both the adaptive origins of hindwing tails and their frequent loss across the phylogeny.

The macroevolutionary lability of blue-green wing coloration may also reflect an evolutionary arms race with visually oriented predators, consistent with the Biotic Challenge hypothesis, which posits strong selection on prey by specialized insectivorous birds (Sherry et al. 2020). In Ithomiini butterflies, for instance, competition within mimicry rings may erode the antipredator effectiveness of aposematic signals (Chazot et al. 2025). In Eudaminae, blue-green coloration was associated with increased speciation rates, yet transitions away from the trait were frequent, suggesting that its adaptive advantage may vary with ecological context. The prevalence of shared warning signals associated with prey evasiveness has been postulated when the costs of evading capture or educating predators are high (Ruxton et al. 2004), although such costs need to be thoroughly quantified before proposing that blue-green iridescence shared by many Eudaminae species is linked to mimicry.

Potential biases and limitations were accounted for during data analyses and the interpretation of results. Although our taxon sampling is incomplete, it is broadly balanced across tribes, and sensitivity analyses excluding widespread species in HiSSE analyses, which substantially reduced statistical power, still recovered the same macroevolutionary patterns.

Transition-rate estimates may also be sensitive to missing taxa. However, congruent patterns recovered from both HiSSE and Markov models in phytools support their robustness. Finally, Ornstein-Uhlenbeck models are prone to overfitting adaptive optima (Cooper et al. 2016), yet explicit comparisons against Brownian Motion models showed strong support for environmental effects on hindwing-tail evolution only. The extensive convergent evolution of antipredator traits in Eudaminae, also documented in previous work and tested *via* simulations and against Brownian Motion models (Linke et al. 2025a), points against neutral drift as the main driver.

Nevertheless, some limitations in trait coding remain. Blue-green iridescence might be difficult to classify consistently into discrete categories, especially when based on photographed specimens, whereas our environmental regime assignments (tropical *vs*. temperate occurrence or diurnal *vs*. non-diurnal activity) may oversimplify ecological variation, such as differences in microclimates or predator assemblages. However, we expect the effects of any such influence on parameter estimates to be minor, given the consistency across analyses.

### Antipredator Defenses Promote Diversification in Specific Environmental Contexts

Our biogeographical and evolutionary transition analyses revealed asymmetric shifts between tropical and extratropical regions. Dispersal into temperate regions was more common than back dispersals to the tropics. Yet, when transitions were modeled jointly with wing traits, we recovered the opposite pattern, mirroring the post-mid-Miocene sudden increase in relative dispersal rates out of temperate regions (Fig. 3b). We found no evidence that temperate breeders have increased speciation rates, consistent with studies showing that geographic range expansion, rather than climatic zone, better predicts diversification rates (Li and Wiens 2022).

Interestingly, transitions toward blue-green wing coloration occurred more frequently in lineages shifting from temperate to tropical regions, explaining the higher prevalence of this trait in extant tropical species of the subtribe Eudamina (Linke et al. 2025a). This pattern aligns with expectations of higher predation pressure in the tropics than in temperate regions (Roslin et al. 2017), where resource partitioning and specialization among insectivorous birds are greater (Sherry et al. 2020), and where intense predation could promote warning coloration (Medina et al. 2025).

Non-diurnal Eudaminae exhibited comparatively low speciation and phenotypic transition rates. Shifts in diel activity alter exposure to predator guilds, potentially involving nocturnal vertebrates or ambush arthropods rather than diurnal birds. If indeed predation pressure is stronger in non-diurnal than in diurnal Eudaminae, reduced speciation rates are consistent with theoretical expectations that strong predation reduces population sizes, decreases interspecific competition, and weakens disruptive selection (Pontarp and Petchey 2018; Chaparro-Pedraza et al. 2022). Contrary to other nocturnal Lepidoptera, hindwing tails are absent in extant non-diurnal Eudaminae, possibly because aerodynamic constraints in strong-flying skippers might render long tails ineffective for powered flight.

### Environment-Dependent Adaptive Evolution of Antipredator Defenses

Hindwings contribute to evasive flight but are not essential for general flight in Lepidoptera (Jantzen and Eisner 2008), while tails influence aerodynamic performance in swallowtail butterflies (Park et al. 2010). However, this remains to be studied in tailed species capable of fast, highly maneuverable flight with high wing-beat frequencies (e.g., Eudaminae or Anaeini leafwing butterflies). Alternatively, Eudaminae tails may offer limited protection against predators that rely primarily on non-visual senses, such as bats. Moth hindwing tails are usually twisted, which improves sonar deflection, whereas tails in butterflies exhibit a non-twisted structure. By contrast, transitions from non-diurnal to diurnal activity often involved the evolution of blue-green coloration (Fig. 4b). This is consistent with potentially mimicry-driven defenses (either Müllerian or Batesian) observed in the genus *Phocides* Hübner, a clade within the predominantly non-diurnal tribe Phocidini that mimics diurnal aposematic firetip skippers (subfamily Pyrrhopyginae). Overall, these results indicate that non-diurnal lineages are relatively evolutionarily stable, but transitions back to diurnality frequently involve the evolution of warning coloration as an antipredator defense.

Our study demonstrates that antipredator defenses shaped by environmental variation have played a key role in driving both phenotypic and species diversification in Eudaminae. Hindwing tails and blue-green wing coloration appear to evolve under strong selection imposed by predators, yet their frequent loss highlights the dynamic nature of these defenses over macroevolutionary timescales. The two traits are likely under different selective pressures, either during predator detection and identification (blue-green coloration) or during predator attack (hindwing tails), and they show largely decoupled evolutionary trajectories. Disruptive selection may favor extreme morphologies or colorations, thereby promoting diversification.

Alternatively, antipredator traits may be associated with traits involved in mate choice, such as body size (Marchinko 2009), which may partly explain the link between blue-green wing coloration and size in Eudaminae (Linke et al. 2025b). Overall, our findings show that predator-mediated selection and environmental variation jointly drive the evolution and repeated loss of antipredator traits, shaping both butterfly phenotypic diversity and lineage diversification.

## Acknowledgments

We are very thankful to Yves Basset, Ranjit Sahoo, Ullasa Kodandaramaiah, and Andrew Warren for sharing specimens/DNA aliquots for this study. We are grateful to Diana Silva and SERFOR for assistance with research permits (Ministerio de Agricultura, Peru; authorization Nos., AUT-IFS-2017-048, AUT-IFS-2021-007, AUT-IFS-2024-79). We thank Blanca Huertas at the Natural History Museum in London and Théo Léger at the Museum für Naturkunde in Berlin for facilitating access to specimens archived in their collections. We acknowledge the computational resources provided by the CESNET LM2015042 and the CERIT Scientific Cloud LM2015085, provided under the programme "Projects of Large Research, Development, and Innovations Infrastructures". We thank the Instituto Chico Mendes de Conservação da Biodiversidade (ICMBio) for Brazilian collection permits 10438-8 and 10802-38. This study is registered under the Brazilian SISGEN (XXXXXX).

## Data Availability

All the original data and scripts necessary to reproduce the analyses reported in this study can be accessed through the Dryad link. License types for the images produced for this study are under the Creative Commons Zero (CC0). Short read sequence data are available on the Sequence Read Archive (SRA) at the NCBI under the BioProject ID XXXXXX. DNA sequence alignments, all data analyses files, and scripts are available on Dryad. Supplementary figures and tables are available online.

## Supplementary Figures and Tables

### Supplementary Tables

**Table S1**: Photographed Eudaminae specimens archived in museums and research institutions (1,349 specimens representing 281 species; median of five specimens per species). Institutional abbreviations: **ACHC**, Academia de Ciencias (Instituto de Ecología y Sistemática), Boyeros, La Habana, Cuba; **BCCAS**, Biology Centre of the Czech Academy of Sciences, Institute of Entomology, České Budějovice, Czechia; **BMNH**, Natural History Museum, London, UK; **MNLB**, Museum für Naturkunde, Leibniz Institüt für Evolution und Biodiversitätsforschung, Berlin, Germany; **MUSM**, Museo de Historia Natural, Universidad Nacional Mayor de San Marcos, Lima, Peru; **BoA**, Illustrated Lists of American Butterflies, (http://www.butterfliesofamerica.com/). Forewing (FW) length, hindwing (HW) length, and hindwing tornus (HW.tornus) were measured in centimeters. Tail ratio is defined as HW.tornus / HW.length. Linear measures correspond to those illustrated in Figure 1 of the main manuscript.

**Table S2**: Trait database for Eudaminae butterflies. For each described species, we report: (**i**) presence (1) or absence (0) of hindwing tails; (**ii**) presence (1) or absence (0) or blue-green iridescent coloration; (**iii**) diel activity, coded as diurnal (0) or non-diurnal (1); and (**iv**) geographic category, distinguishing tropical breeders, non-tropical breeders, and widespread species occurring as occasional strays in temperate regions. Uncertain values were assigned a question mark “?”.

**Table S3**: Specimens used to infer molecular phylogenies, including 281 Eudaminae species and 29 Hesperiidae outgroups. Locality information and the institutions where specimens were deposited are provided. For samples sequenced in previous studies, we list the sequencing method: **WGS**, whole-genome sequencing, **ahe_1.0**, target enrichment using the BUTTERFLY probe kit 1.0 (Espeland et al. 2018), **ahe_2.0**, BUTTERFLY probe kit 2.0 (Kawahara et al. 2018). Accession numbers refer to NCBI BioProject, BioSample or SRA entries. The number of loci recovered for this study is reported for each specimen.

**Table S4**: Biogeographical and trait data for species included in the molecular phylogeny. Linear morphological measurements (FW.length, HW.length, HW.tornus; in centimetres) are per-species averages derived from the specimen measurements listed in Table S1. Phylogenetically corrected tail length (rel.tail) represents the residuals from a log-log phylogenetic regression of HW.tornus on FW.length and HW.length. Wing-trait data are sourced from the Eudaminae trait database (Table S2). Species geographic ranges were assigned to seven bioregions: 1) Mesoamerica and the Pacific slope of the Andes, 2) the Antilles, 3) the Andes above ∼1,000 m, 4) Amazonia and the eastern foothills of the Andes, and 5) the Atlantic Forest, and two extratropical regions, 6) the Nearctic and 7) the Pampas.

**Table S5**: Fitted Ornstein-Uhlenbeck (OU) and Brownian motion (BM) models evaluating the influence of three environmental regimes on the evolution of hindwing tail length in Eudaminae: (1) tropical diurnal, (2) tropical non-diurnal, and (3) temperate diurnal (all temperate species are diurnal). Six evolutionary models were compared: (1) **OU1**, a single trait optimum (θ) and shared evolutionary rate (σ^2^) and selection strength (α) across regimes; (2) **OUM**, distinct θ per regime; (3) **OUMV**, distinct θ and σ^2^; (4) **OUMA**, distinct θ and α; (5) **BM**, single-rate Brownian motion model; and (6) **BMS**, Brownian motion with regime-specific σ^2^. Model comparison using small-sample corrected Akaike weights (weighted AICc) supported the **OUMV**, indicating that hindwing tails evolve toward different optima under each environmental regime and with different evolutionary rates. Under this model, tropical diurnal species exhibit longer hindwing tail length than tropical non-diurnal and temperate species. Parameter estimates represent averages across 100 stochastic character mappings. Analyses were repeated using both the MCMCTree and BEAST trees.

**Table S6**: Fitted Ornstein-Uhlenbeck (OU) and Brownian motion (BM) models evaluating the influence of blue-green iridescent coloration on wings on the evolution of hindwing tail length in Eudaminae. Six evolutionary models were compared: (1) **OU1**, a single trait optimum (θ) and shared evolutionary rate (σ^2^) and selection strength (α) across regimes; (2) **OUM**, distinct θ per regime; (3) **OUMV**, distinct θ and σ^2^; (4) **OUMA**, distinct θ and α; (5) **BM**, single-rate Brownian motion model; and (6) **BMS**, Brownian motion with regime-specific σ^2^. Model comparison using small-sample corrected Akaike weights (weighted AICc) supported the **BMS**, indicating that hindwing tails do not evolve toward different adaptive optima under the presence of iridescent wing coloration, but probably have different evolutionary rates. Parameter estimates represent averages across 100 stochastic character mappings. Analyses were repeated using both the MCMCTree and BEAST trees.

**Table S7**: Speciation rate estimates under the Hidden State-dependent Speciation and Extinction (HiSSE) framework implemented in RevBayes. Four discrete states combine environmental and phenotypic attributes: TDO, tropical diurnal species lacking the focal trait (hindwing tails or blue-green coloration); TNO, tropical non-diurnal species lacking the trait; TDW, tropical diurnal species expressing the trait; CDO, temperate diurnal species lacking the trait. Pairwise differences in speciation rates (λ) summarized with 95% credible intervals. Comparisons whose intervals do not overlap zero are marked with an asterisk and in bold. Differences are reported as state_1_ state_2_, so positive values indicate higher speciation rates in the state 1. Posterior probabxilities represent the proportion of posterior samples in which the speciation rate of one state exceeded that of the other. The dataset is reduced compared to the analyses presented in Table 2 in the main manuscript, in that here tropical species with stray into temperate regions (i.e., widespread) were removed from the analyses.

## Supplementary Figures

**Figure S1**: Time-calibrated phylogeny of Eudaminae inferred in BEAST using 70 loci selected for clock-likeliness and phylogenetic information content (SortaDate) alongside 14 widely used legacy loci in butterfly systematics. Branches are labelled with posterior probabilities, and nodes show 95% highest posterior density (HPD) intervals. Scale is in millions of years.

**Figure S2**: Seven competing models of discrete trait evolution were evaluated using the fitMK function in phytools: 1) full model, all transition rates free to vary; 2) symmetric model, forward and backward transitions constrained to be equal; 3) equal-rates model, all transition rates equal; 4) no simultaneous shifts in wing phenotype expressing antipredator defense and diel activity (no transitions between TDW and TNO); 5) no simultaneous shifts in antipredator wing phenotype, diel activity and geographic region (no transitions between TDW and TNO or between TDW and CDO); 6) no simultaneous shifts in diel activity, antipredator wing phenotype and geographic region (no transitions between TNO and TDW or between TNO and CDO); and 7) transitions restricted to the tropical diurnal state lacking antipredator phenotype (TDO). Transition rates between states under the best-fit models: restricted transitions for hindwing tails (only involving the tropical diurnal state and the tailless phenotype; **model 7**) and all-rates-different for blue-green coloration (**model 1**). The number of species in each state is indicated as proportional gray circles.

**Figure S3**: Molecular phylogeny of Eudaminae inferred in IQ-TREE using 369 loci with >70% species coverage. Branches are labelled with ultrafast bootstrap (**UFBoot**) followed by Shimodaira-Hasegawa-like approximate likelihood-ratio test (**SH-aLRT**) values. Branch lengths and the scale bar represent substitutions per site.

**Figure S4**: Molecular phylogeny of Eudaminae inferred in ExaBayes using 369 loci with >70% species coverage. Branches are labelled with posterior probability values, and branch lengths and the scale bar represent expected substitutions per site.

**Figure S5**: Species tree of Eudaminae inferred in wASTRAL using 369 gene trees, with taxa having >50% missing data removed. Branches are labelled with local posterior probability values, and branch lengths and the scale bar are in coalescent units.

**Figure S6**: Time-calibrated phylogeny of Eudaminae inferred in BEAST using 70 loci with <50% missing data and >75% species coverage, alongside 14 widely used legacy loci in butterfly systematics. Branches are labelled with posterior probabilities, and nodes show 95% highest posterior density (HPD) intervals. Scale is in millions of years.

**Figure S7**: Time-calibrated phylogeny of Eudaminae inferred in MCMCTree using 369 loci with >70% species coverage, with the IQ-TREE topology as a fixed guide tree. Nodes show 95% highest posterior density (HPD) intervals. Scale is in 10 million years (1 unit = 10 Myr).

**Figure S8**: Ancestral biogeographic ranges of Eudaminae inferred under the DEC model in BioGeoBEARS. Extant species were assigned to seven bioregions: five tropical regions, **(M)** Mesoamerica and the Pacific slope of the Andes, **(C)** the Antilles, **(A)** the Andes above ∼1,000 m, **(Z)** Amazonia and the eastern foothills of the Andes, and **(F)** the Atlantic Forest, as well as two extratropical regions, **(N)** the Nearctic and **(D)** the Pampas (Table S4). The Asian genus *Lobocla*, representing a single dispersal event out of the Americas, was excluded from the analyses. The first phylogeny shows the most probable range at each node, while the second phylogeny depicts the probabilities of alternative ranges at each node.

**Figure S9**: Estimated dispersal events among biogeographical regions inferred under the DEC model using 5,000 Biogeographical Stochastic Mapping replicates in BioGeoBEARS. Arrows indicate the direction and number of inferred dispersal events. Only connections with more than two events are shown. Node layout reflects the strength of biogeographical interconnections, arranged using the force-embedded algorithm in the R package qgraph. Biotic interchange through time was estimated in three time bins: (1) between the most recent common ancestor of Eudaminae in the mid-Eocene and the mid-Miocene (44–15 Myr), (2) mid-Miocene to early Pliocene (15–5 Myr), and (3) early Pliocene to the present (5–0 Myr).

