## Supplementary figures and images for "Environmental Variation Promotes Convergent Evolution and Rapid Diversification of Wing Shape and Color in Skipper Butterflies"

### Figure S1

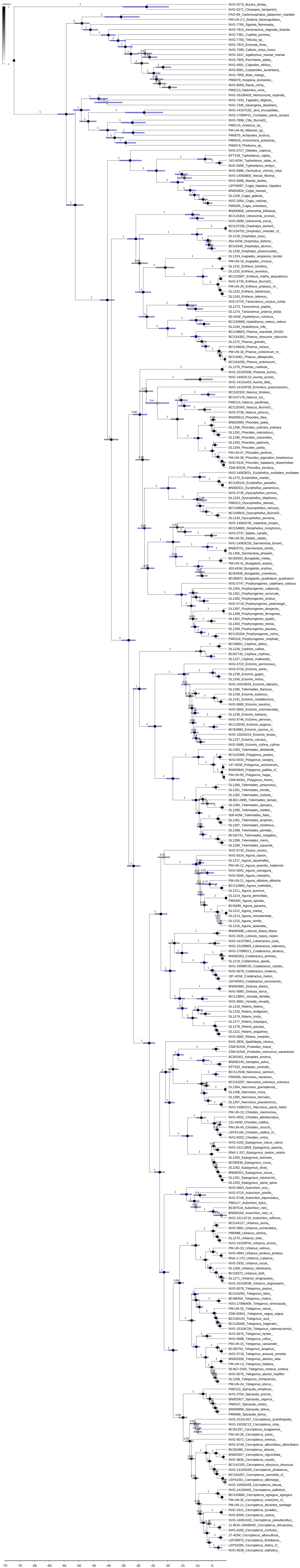

### Figure S2

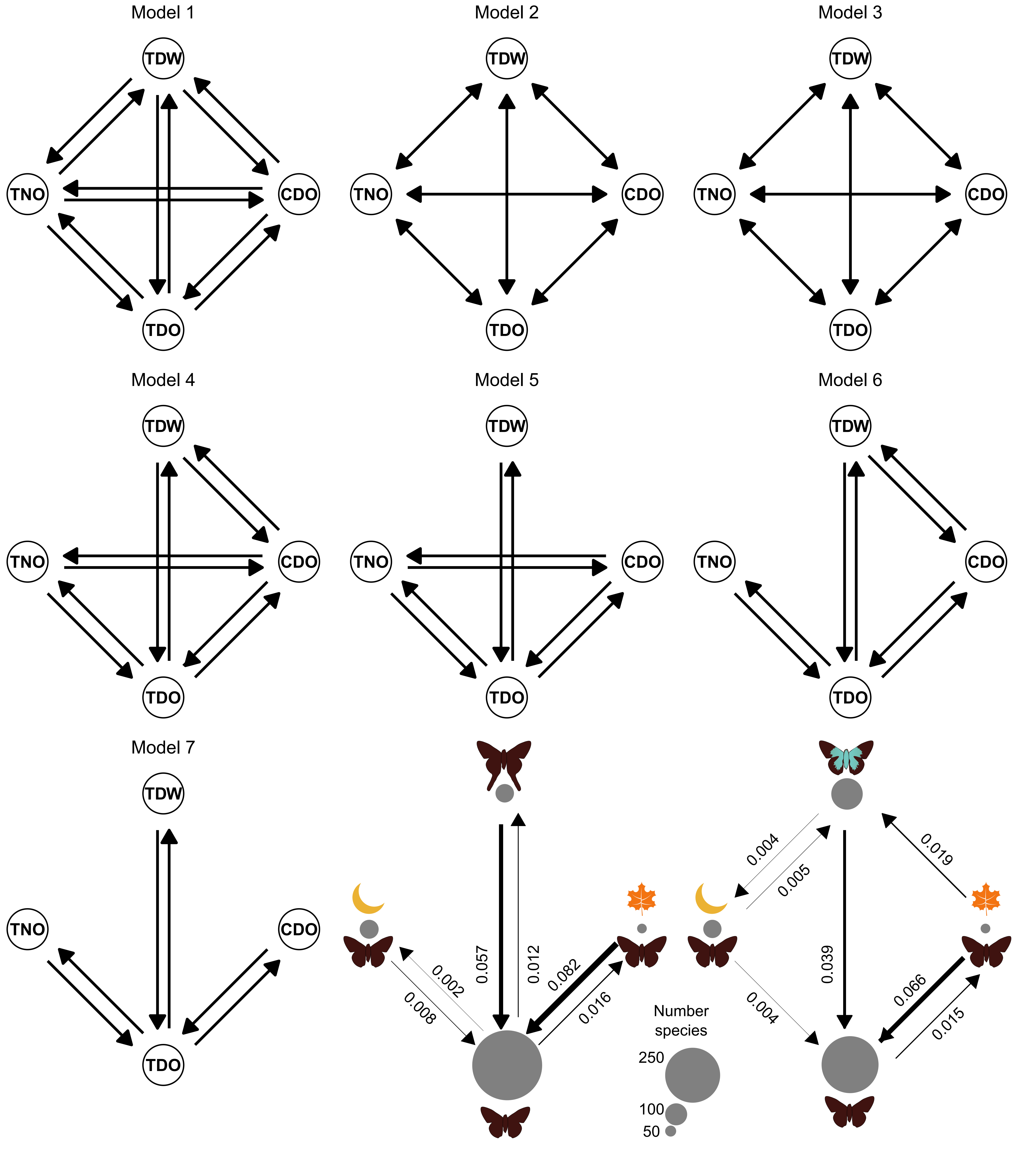

### Figure S3

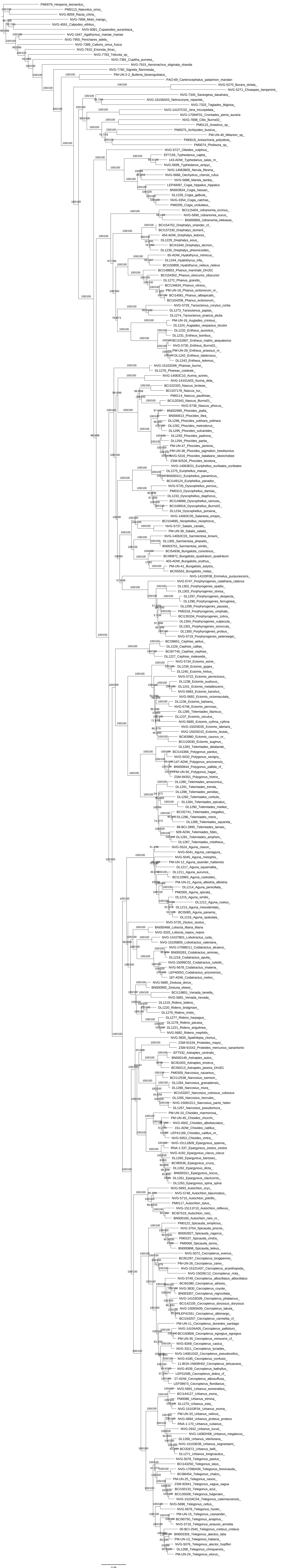

### Figure S4

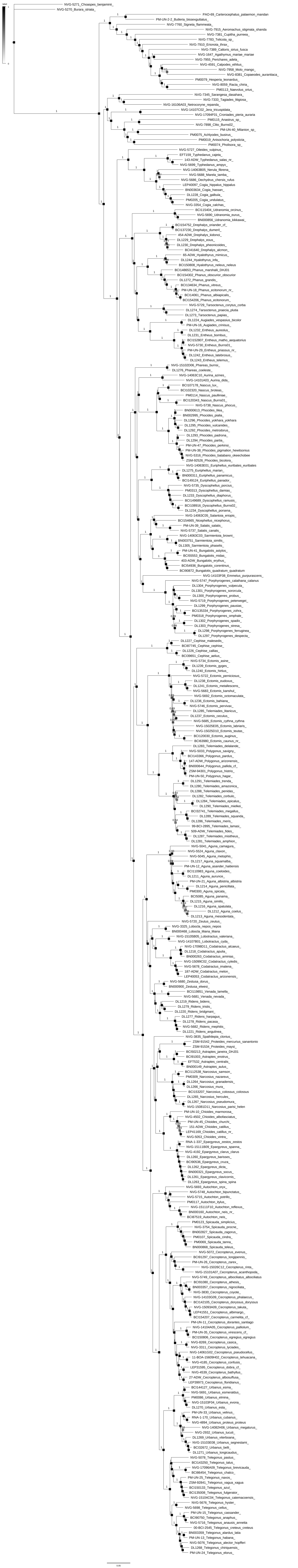

### Figure S5

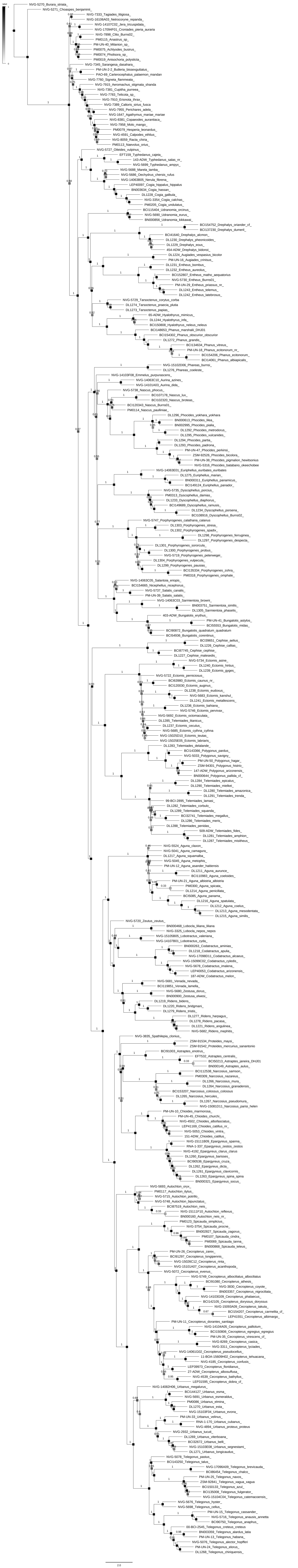

### Figure S6

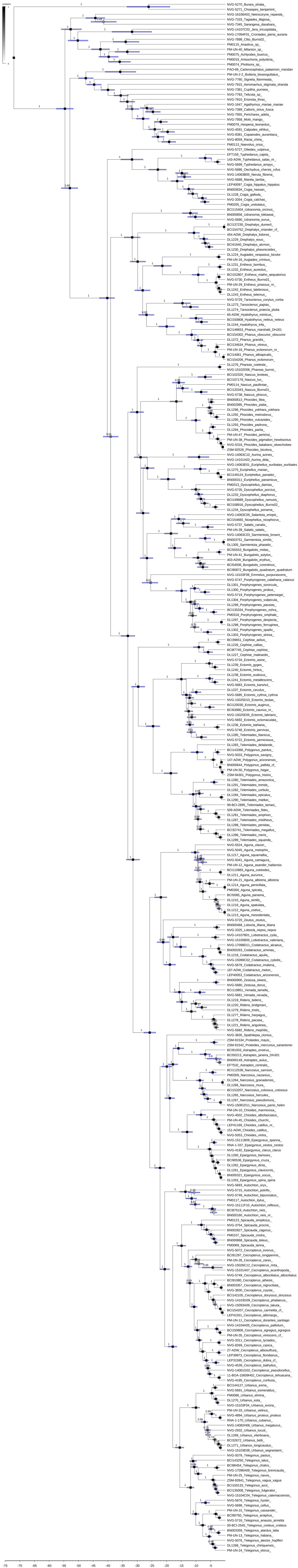

### Figure S7

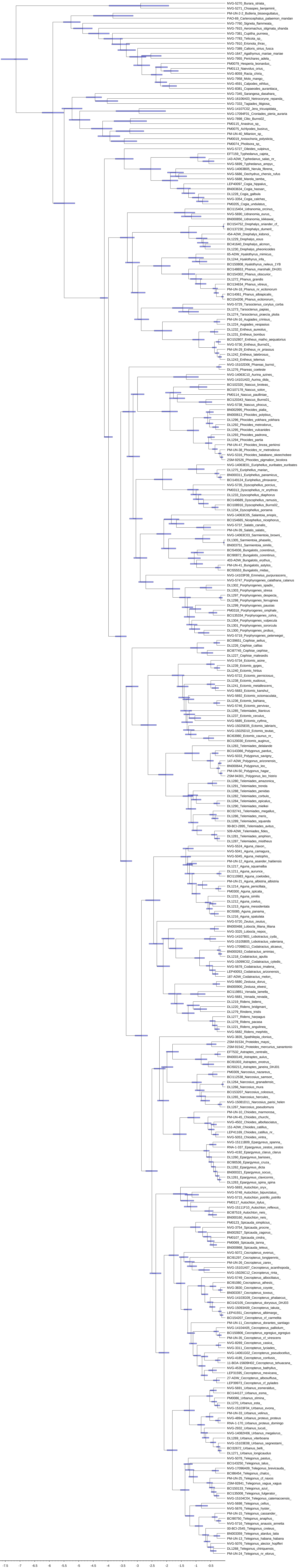

### Figure S8

**BioGeoBEARS DEC on eudaminae unconstrained**
**ancstates: global optim, 7 areas max. d=0.0219; e=0.0011; j=0; LnL=−1100.83**

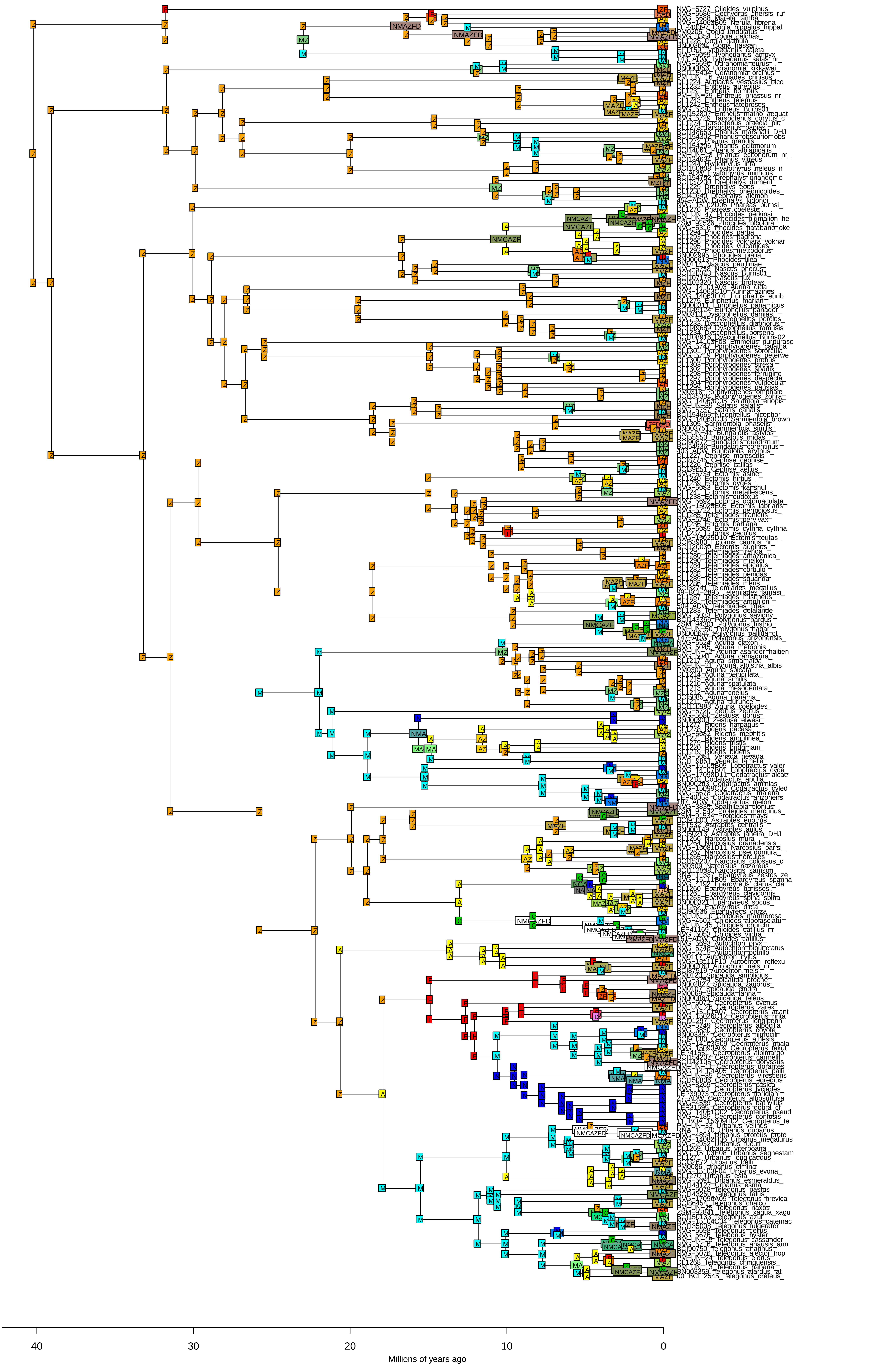

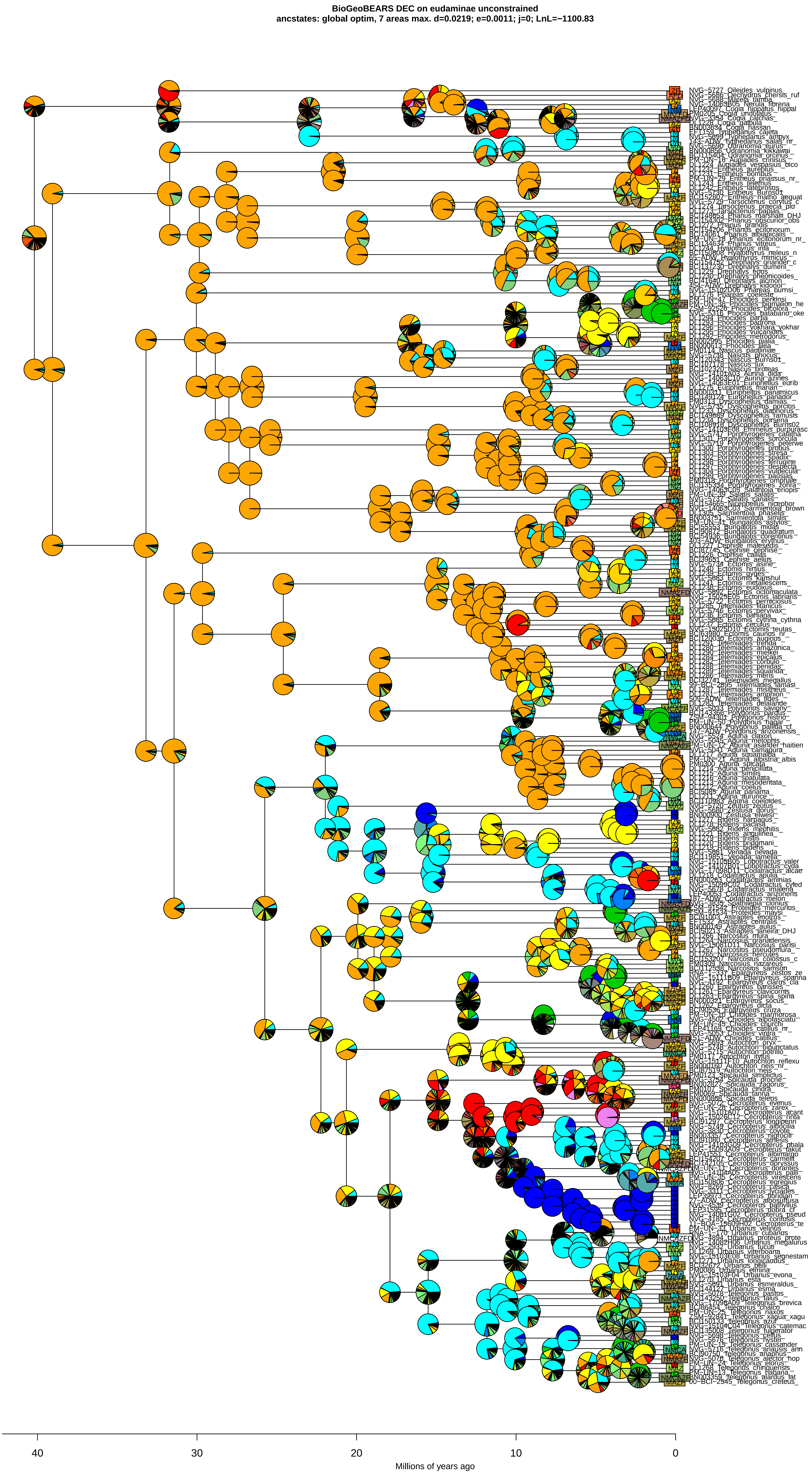

### Figure S9

Total Dispersals

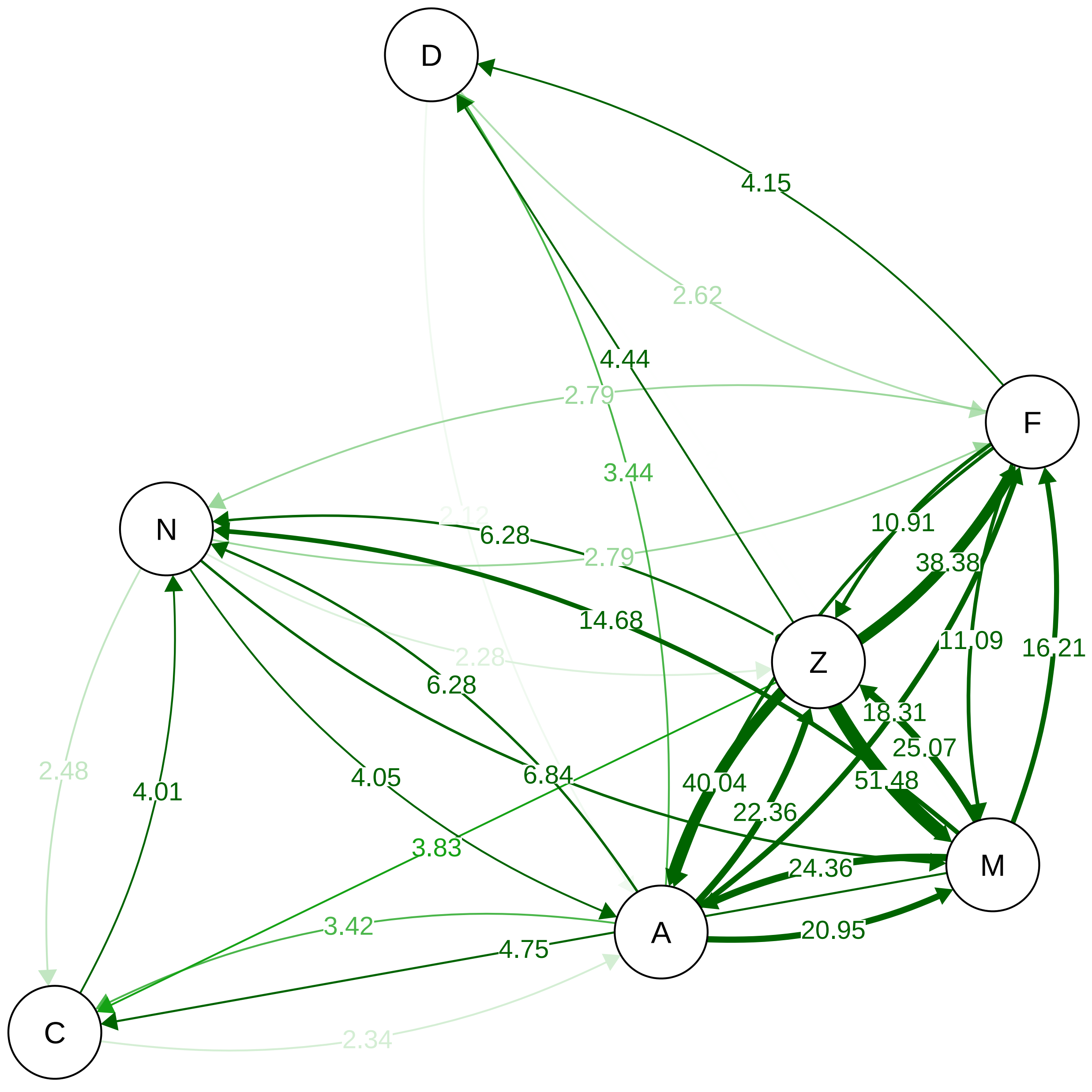


Dispersals 0−5 Myr

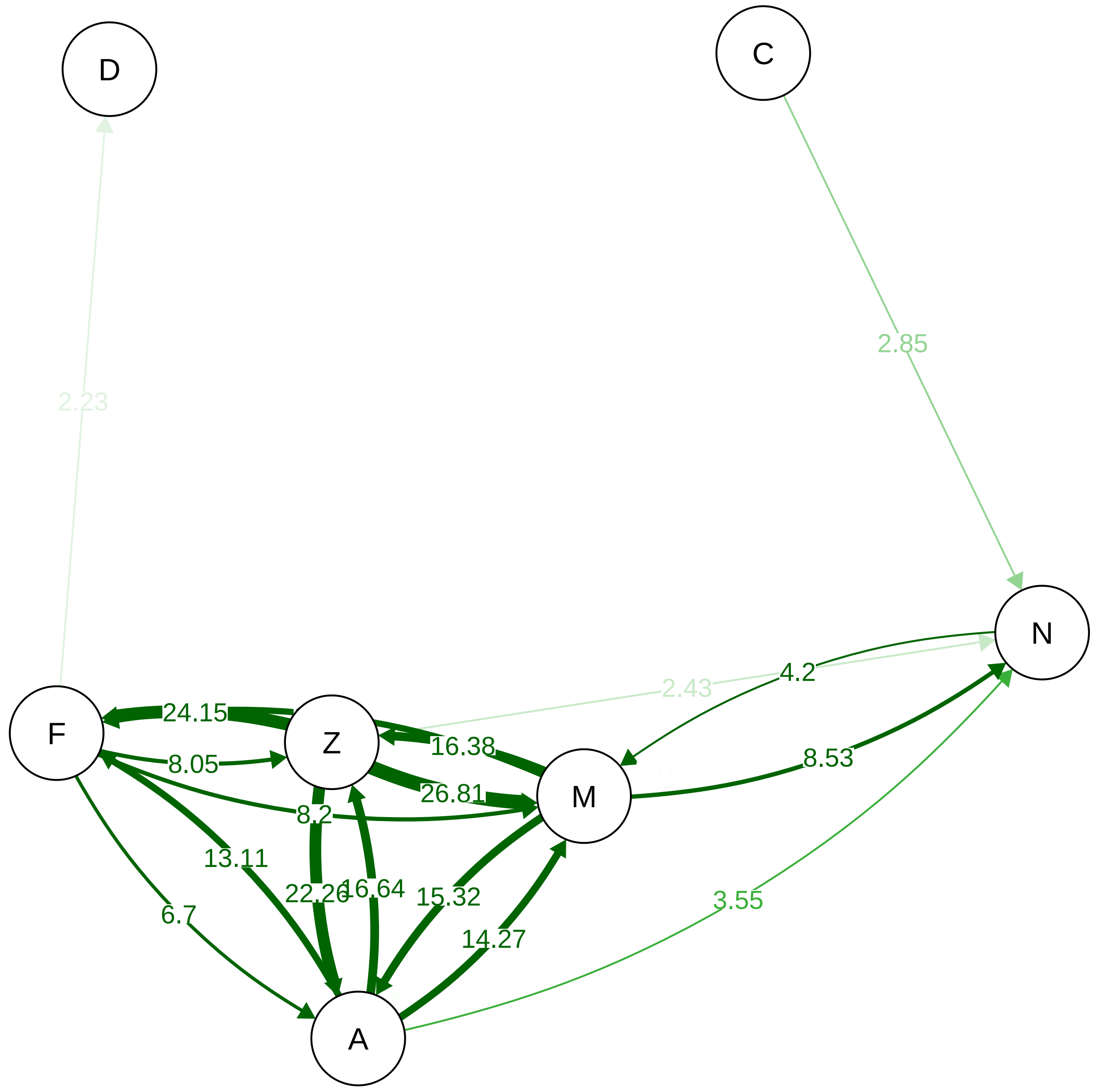


Dispersals 5−15 Myr

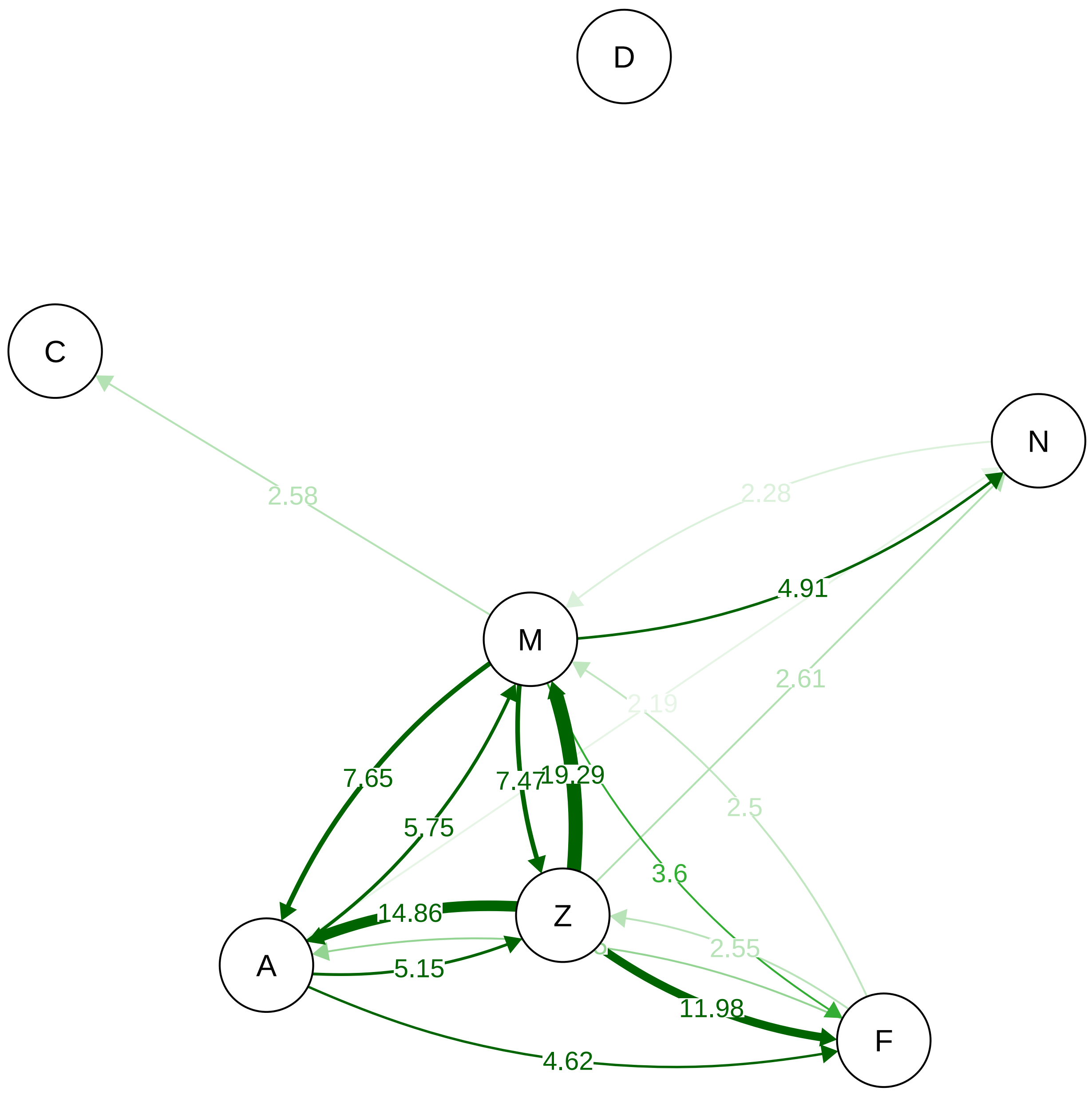


Dispersals 15−44 Myr

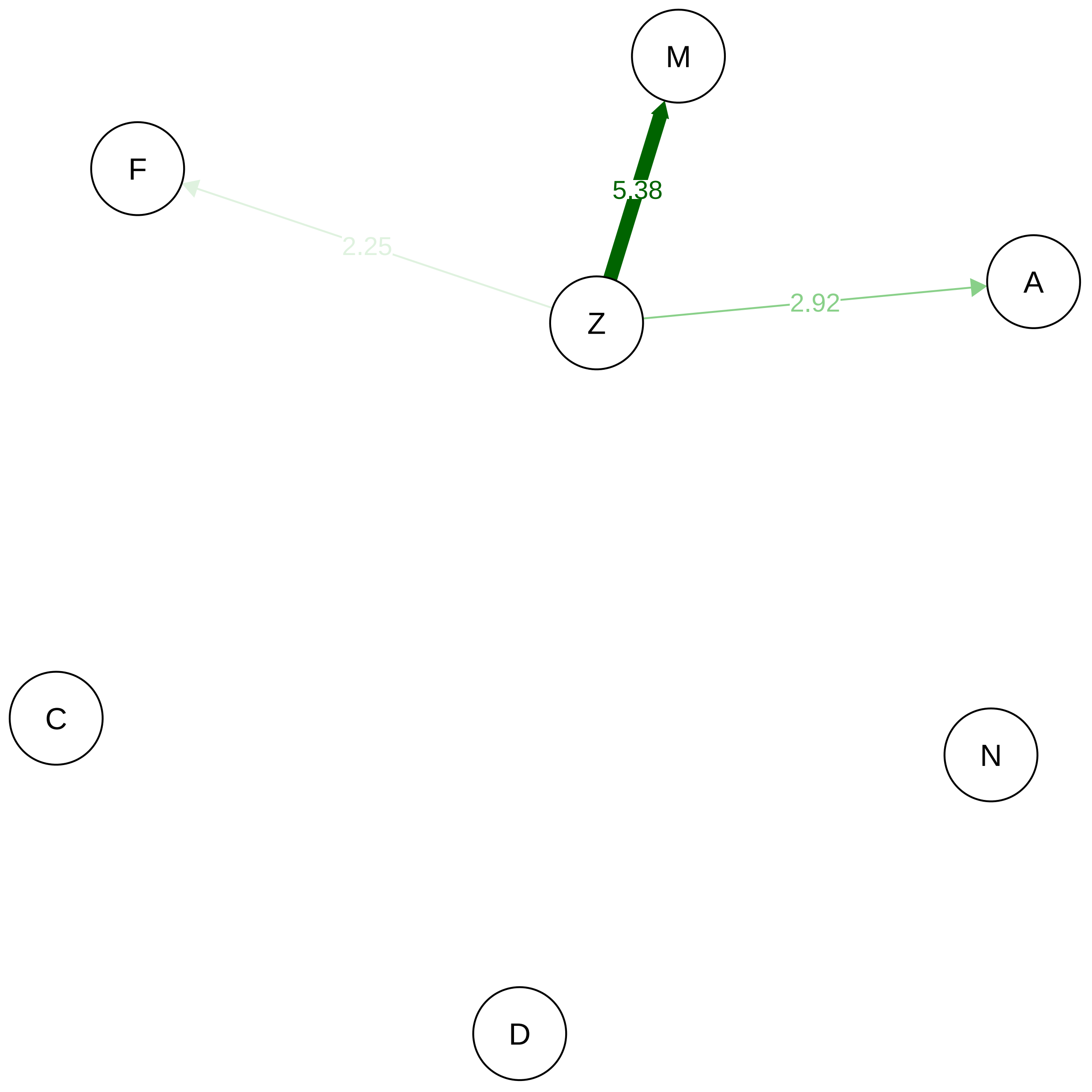
